# Salicaceae as potential host plants of *Xylella fastidiosa* in European temperate regions

**DOI:** 10.1101/2022.06.10.495618

**Authors:** Noemi Casarin, Séverine Hasbroucq, Lena Pesenti, Amandine Géradin, Amélie Emond, Júlia López-Mercadal, Miguel Ángel Miranda, Jean-Claude Grégoire, Claude Bragard

## Abstract

The discovery of three subspecies of *Xylella fastidiosa* in Europe has triggered major attention on the potential spread up North of the bacteria. Assessing the susceptibility of a previously unexposed European flora is a key element that remains widely unknown. Under biosafety facility, we evaluated the susceptibility of Salicaceae such as *Populus tremula, Populus canescens, Salix alba* and *Salix caprea* by mechanically inoculating the KLN59.3 GFP-labelled *X. fastidiosa* at 22 °C and at 28 °C. Bacterial movement and multiplication in plants were investigated by PCR, real-time PCR, confocal or scanning electron microscopy. Nine months post-inoculation, 100 % of the plants tested positive for *X. fastidiosa*, with the exception of 57% for *P. canescens* under the 22 °C-growing conditions. Bacteria were detected up to 120 cm from the inoculation point for *S. alba*. They were detected in the roots of all species and were successfully isolated for *S. alba* and *P. tremula*. Estimates of average CFU/g of plant tissue per species ranged from 1.5E + 03 to 3.5E + 06, with the lowest figures for *P. canescens* and the highest for *P. tremula* together with high number of totally obstructed vessels observed by confocal microscopy. The possibility of insect transmission was also evaluated using an experimental set up based on Mallorca Island. There, transmission by *P. spumarius* of both *X. fastidiosa* ST1 and ST81 was proven on *S. alba*. We thus demonstrated that indigenous European Salicaceae such as *S. alba* or *P. tremula* are new potential hosts for *X. fastidiosa*.

## 1 INTRODUCTION

The American-native phytopathogenic bacterium *Xylella fastidiosa* (Xanthomonadaceae; Wells et al., 1987) has recently shaken Europe. First identified on olive trees in Southern Italy in 2013 (EFSA, 2013; Saponari et al., 2013; Martelli et al., 2016), it has led to major economic, environmental and social damage (FAO IPPC, 2017; EFSA, 2018). Later on, it was also reported in Southern France, Corsica, Continental Spain, Balearic Islands, and in Portugal on a variety of hosts (EFSA, 2021). Divided into several subspecies, including subsp. *fastidiosa*, subsp. *multiplex* and subsp. *pauca* (Schaad et al., 2004) and into more than 87 sequence types (ST), the diversity of the detected European strains suggests multiple independent introductions on the continent (EFSA, 2018; Landa et al., 2020). The numerous interceptions of *X. fastidiosa* in traded goods at border controls (EFSA, 2018; EUROPHYT online) confirm that entry events of the bacterium are not scarce. Once introduced, *X. fastidiosa*, strictly limited to the xylem vessels of plants and to the foregut of its xylem sap-feeding insect vectors (Chatterjee et al., 2008), colonizes plants and spreads in new environments following a complex pattern defined by the interactions of all the components of its pathosystem (EFSA, 2015). The *Xylella*-pathosystems are known to be highly polymorphic leading to different diseases in various regions, each one having their own range of host plants, insect vectors, bacterial strains and environmental conditions.

As shown by the current European outbreaks, the threat for Mediterranean areas is definite, with efficient hosts-vectors-strains interactions and particularly favorable climatic conditions for *X. fastidiosa* development (Purcell, 1997; Bosso et al., 2016; Sicard et al., 2018). On the other hand, the potential impact in northern European regions outside of its present distribution range, remains uncertain (Occhibove et al., 2020). Despite *X. fastidiosa* has been generally associated to warm areas (i.e., Mediterranean climate) (Bosso et al., 2016), temperate climates do not seem to be a barrier for bacterial growth, as some strains have already been encountered in colder areas e.g. Canada and North-Eastern USA (Goodwin and Zhang, 1997; Zhang et al., 2011; Guan et al., 2014). In addition, prediction models based on species distribution model (SDM) techniques showed that northern and eastern regions of Europe, including Belgium, the Netherlands and the coastal regions of UK and Ireland, have suitable climates for a potential bacterial establishment (EFSA, 2019; Godefroid et al., 2019). Therefore, the risk of an establishment for these regions depends mostly on the presence of efficient hosts and vectors combinations. These combinations are challenging to identify because of pathosystem complexity and polymorphism, and extrapolating from current knowledge of native area has proven not to be sufficient to predict new outbreaks, as illustrated by the Olive Quick Decline Syndrome (OQDS) in Italy (Krugner et al., 2014). On the insect side, xylem-feeding specialists, mainly spittlebugs and leafhoppers (Hemiptera: Cicadomorpha), were reported throughout all the European continent (de Jong, 2013; EFSA, 2015), making it possible for the bacteria to spread up north. Regarding plants, the molecular determinants of the host range of *X. fastidiosa* are far from understood despite the availability of several genomes sequenced (Sicard et al., 2018; Denancé et al., 2019; Nunney et al., 2019). The host plant database compiled by EFSA (2022) is extensive, comprising up to 655 species belonging to 88 families and continues to grow with the discovery of new hosts, mainly through systematic sampling of plants in natural environments in Europe. This constant identification of new hosts emphasizes that many of them are still unknown, as they had never been tested or exposed to the bacterium, given the differences of flora from *Xylella*-native areas.

Sustained investigation of the host range, and testing the efficiency of new potential hosts, is therefore essential in northern Europe to determine where to focus resources and increase the probabilities of early detection. The testing possibilities being wide, one way to start is to investigate plant species associated with potential insect vectors, drivers of bacterial spread. The Salicaceae (*Salix* spp. and *Populus* spp.), widely distributed in the northern hemisphere, have been associated with several potential vectors for Europe, including *Philaenus spumarius, Aphrophora alni, Aphrophora salicina* and *Cercopis vulnerata* (Nickel and Remane, 2002; Nickel, 2008; Hasbroucq et al., 2020). Furthermore, adapted to humid soil, this plant family is abundant in riparian areas, which have been pointed out in America as a favorable niche for both *X. fastidiosa* and its vectors (Purcell, 1974; Purcell, 1975; Purcell and Saunders, 1999). Purcell and Saunders (1999) have detected *X. fastidiosa* in two willow species in Californian riparian areas, *Salix laevigata* (red willow) and *Salix lasiolepis* (arroyo willow), and in one poplar species *Populus fremontii* (Fremont cottonwood), all native to America. Two other surveys in riparian vegetation in California also reported positive *Salix* spp. around vineyards (Freitag, 1951; Costello et al., 2017). However, in all the surveys, the description of symptoms is missing. Nevertheless, hosts could also remain asymptomatic and act as a reservoir in the environment, the bacterium behaving as an endophyte (EFSA, 2018).

In this study, European riparian Salicaceae species have been hypothesized as potential hosts and natural reservoirs for *X. fastidiosa* in European maritime climate areas. We monitored bacterial spread and multiplication in the plants after mechanical inoculations of *X. fastidiosa* subsp. *fastidiosa* GFP-labeled strain KLN59.3 (Newman et al., 2003). The GFP was used to investigate the distribution of bacteria in xylem vessels. Scanning electron microscopy was also performed along with real-time quantitative PCR and re-isolation of the bacteria. In parallel, in the Balearic Islands, transmission tests with naturally infected *Philaenus spumarius* collected in the field were also carried out on young seedlings of *Salix alba* and *Populus tremula*.

## 2 MATERIALS AND METHODS

### 2.1 Collection of plant material

Two poplar and two willow species were used for different experiments: *Populus tremula, Populus canescens, Salix alba* and *Salix caprea*. Plants of *P. tremula* (1+0 Vloethemveld origin) were bought at the Sylva nursery in Ghent (Belgium). *Salix alba* cuttings were made from plants (90/120 Salix alba “Liempde”) bought at the Lillois nursery in Lillois-Witterzée. Cuttings of *P. canescens* were made from roots of *P. canescens* trees near Louvain-la-Neuve and cuttings of *S. caprea* from the branches of trees located in the botanical garden Jardin Massart in Brussels. The plants were potted and were set in an insect-proof greenhouse, until the shoots were 40-60 cm high and strong enough to stand mechanical inoculation. Tobacco plants (*Nicotiana tabacum*) cultivar “Petit Havana SR1” and periwinkle (*Catharanthus roseus*) “Pacifica XP Polka dot” were used as control plants to verify inoculation success. No fertilizer was brought to the plants during the whole experiment. Plants were tested by PCR before inoculation trials to verify previous absence of *X. fastidiosa*.

### 2.2 Preparation of *X. fastidiosa* KLN59.3 inoculum

The *X. fastidiosa* subsp. *fastidiosa* KLN59.3 strain, expressing the Green Fluorescent Protein (GFP), was provided by Professor Steven E. Lindow (UCBerkeley). For the inoculation trial, bacteria were grown from -80 °C glycerol storage on BCYE media (Wells et al., 1981) for 8-10 days at 28 °C in a quarantine facility. Then, they were sub-cultured for 10 more days on Buffered charcoal yeast extract (BCYE) agar at 28 °C. The inoculum was prepared by scraping the 10-day-old colonies from plates and suspending them in Succinate-citrate-phosphate buffer (SCP, pH 7.0; “PM 7/24 (4) *Xylella fastidiosa*”, EPPO 2019) until obtaining a turbid cell suspension. The aliquots were plated and incubated at 28 °C for two weeks to estimate the concentration of the inoculated suspension by colonies counting, which was 1.84 × 10^8^ CFU/ml (DO_600_ : 0.31).

### 2.3 Mechanical inoculation

The plants were not watered three days preceding and after the inoculation to favor the uptake of the bacterial suspension into the xylem vessels. One or two branches per plant were inoculated using the pinprick inoculation method (Hill and Purcell, 1995; Almeida et al., 2001). The basal stem of each branch was pin-pricked at three spots, differently orientated and separated from each other along the stem by 1 cm. One 10 µl drop of inoculum was placed at each spot and pin-pricked 3-4 times with a 26 G sterile needle. During the inoculation, the plants were maintained horizontally for 10-15 min to allow the penetration of the inoculum into the stem. One batch was kept in a temperate quarantine greenhouse (22 °C day, 20 °C night, humidity: 80 %) and another one in a tropical quarantine greenhouse (28 °C day, 25 °C night, humidity: 70 %). *Nicotiana tabacum* and *C. roseus* have been used as control plants to verify the inoculation method. In total, 10 to 13 *P. tremula, P. canescens* and *S. alba*, 6 *S. caprea*, 5 *N. tabacum* and 3 *C. roseus* were inoculated for each temperature condition. Three plants of each species per treatment were also mock-inoculated with SCP buffer only and three additional plants without any treatment were set in each greenhouse. All plants were watered once every two days. Most of them were inoculated in June 2020. However, a small batch of plants was inoculated in 2019 in a preliminary test, while *S. caprea* were inoculated later in March 2021.

### 2.4 Monitoring plant health and bacterial colonization through time

#### 2.4.1 Bacterial detection in petioles with PCR

To follow the progression of the bacteria in the xylem vessels without destructing the stem, the main vein of the leaves and the petioles attached to the inoculated branch were collected at 1, 3 and 6 months post-inoculation (mpi) to be analyzed for bacterial presence. The DNA from plant tissues was extracted following a standard CTAB-based procedure following the requirements of EPPO standard protocols (“PM 7/24 (4) *Xylella fastidiosa*”, EPPO 2019) and then, was tested by PCR of Minsavage et al. (1994). The sampling started from the closest petiole to the inoculation point (IP) and if *X. fastidiosa* was detected, another petiole about 5 cm above was collected and tested. Petiole collection and testing continued until one tested negative to the bacterium. The distance between the IP and the last positive petiole was measured as the current distance of the progression of the bacterium.

#### 2.4.2 Symptom monitoring

The symptoms were monitored once every two weeks. The appearance of wilting, shoot dieback, desiccation, defoliation or any change in leave color were reported and were compared to the control group of plants. The symptoms recorded were then ranked into several categories according to their severity. The size of the plants was measured as well.

### 2.5 Destructive analyses of the plant

At 8-9 mpi, seven plants per species and conditions were destructed to perform analyses on the stems, and on the roots for 1-3 plants per group. The other plants were maintained until 17 mpi to monitor the evolution of their health state on a longer period.

#### 2.5.1 Real-time quantitative PCR

The stem was chopped into 12-15 cm-length segments to assess the bacterial concentration at different levels of the plant and assess the spread of the bacteria. The inoculated branch was divided into three to six pieces depending on the size of the branch. For some plants (2-4 depending on the species), the main stem below the inoculated branch, as well as the roots, were also sampled and divided in similar segments. These samples were cut into smaller pieces and were mashed with a hammer in extraction bags (BIOREBA, Switzerland). The DNA of each sample was then extracted following the standard CTAB-based procedure (“PM 7/24 (4) *Xylella fastidiosa*”, EPPO 2019) and tested by real-time PCR of Harper et al. (2010) targeting bacterial DNA. The bacterial population, represented by estimated log cells/g of plant tissue, was quantified interpolating the Cycle threshold (Ct) obtained by real-time PCR on a linear standard curve (y = -3.8729 + 44.623, R2 = 0.9997) generated using 10-fold serial dilutions of a bacterial suspension with known concentration (from 10^8^ CFU/ml to 10^2^ CFU/ml). Reactions were duplicated and all plates had negative and positive controls. The cut-off values of Ct were set at 37 according to the detection threshold obtained with the 10-fold serial dilutions of *X. fastidiosa* samples.

#### 2.5.2 Re-isolation on BCYE

Re-isolation of the bacterium was performed from a 2 cm-long piece of the stem, at the IP and near the apex of the inoculated branch. The surface of the section was sterilized with bleach (0.5 %, 5 min) and ethanol (70 %, 2 min) and then was rinsed in three baths of sterile water. The segments were peeled with a sterile blade to remove bark and then ground in a BIOREBA bag with a hammer together with 2 ml of SCP buffer. The dilutions were made from the extract and were plated on BCYE media. The plates were incubated at 28 °C and were regularly checked for *Xylella*-like colonies. If present, they were collected and streaked on new BCYE media and incubated again at 28 °C. The isolates were then suspended in biomolecular water and were heated at 100 °C during 5 min. The suspension was tested by PCR of Minsavage et al. (1994) and finally, the amplicons were sequenced to confirm the bacterium identify.

#### 2.5.3 Confocal microscopy

The visualization of the bacterium through confocal microscopy was performed as a result of the expression of the GFP by the mutant strain, as described by Newman et al. (2003). Transversal and longitudinal sections of approximately 55 µm were made with a Leica microtome (SM2000R) from all the segments that were submitted afterwards to real-time PCR, to visualize the distribution of the bacteria along the stem and in the roots. The fine cuts were directly placed on a microscope slide in 50 % glycerol and were observed immediately with a Zeiss confocal microscope (LSM 710) equipped with water objective at magnifications of x 40. The GFP was excited with an argon laser at 488 nm wavelength and observed with a filter selecting the emission spectrum from 493 to 553 nm. High-resolution images were also taken with the Airyscan mode. The visualization was first performed on periwinkles to verify the method and adjust the parameters.

#### 2.5.4 Scanning electron microscopy (SEM)

Longitudinal and cross-sections up to 5 mm thick were made with a Leica microtome (SM2000R) or with a razor blade. They were fixed in 4 % glutaraldehyde diluted in PBS for 24 to 48 hours and were then dehydrated in successive 30 min ethanol baths: 30 %, 50 %, 60 %, 70 %, 90 %, 100 %. The sections were then metalized with a layer of approximately 15 nm of gold deposited using a Cressington Sputter Coater (208 HR) metalliser. They were visualized using a JEOL electron microscope (JBM-7800F) at magnifications ranging from 75 to 3000 times in low magnification (LM), scanning electron microscopy (SEM) and gentle beam (GB-H) modes. The acceleration voltage was fixed at 5 kV and the probe current at 8.

### 2.6 Insect-transmission experiments in Mallorca

Twenty seedlings of *S. alba* and twenty *P. tremula*, equipped with phytosanitary passports and originating from the same Belgian nurseries as for the inoculation trials, were sent to the University of Balearic Islands (UIB) in Palma (Mallorca) in February 2020 and placed in the biosecurity greenhouse at the UIB Campus. Leaves and parts of twigs were collected before the beginning of the transmission experiments to test if the plants were free of *X. fastidiosa*. The transmission experiment was performed twice, in October 2020 and in June 2021, on two different branches of each plant. In October 2020, in total 200 *P. spumarius* adults were collected from infected areas of Mallorca on shrubs, olive and almond trees in the municipalities of Inca and Manacor. Five insects were placed on one netted-branch per plant, on a 20 cm-long section. The insects were kept on the respective branch for a transmission period of four days (approx. 96 hours). After this period, they were collected back and the DNA of their mouthparts was extracted following the standard CTAB-based procedure (“PM 7/24 (4) *Xylella fastidiosa*”, EPPO 2019) and tested for the presence of the bacterium by real-time PCR of Harper et al. (2010). If a Ct < 38 was obtained, the DNA of the respective insect was tested by multilocus sequence typing (MLST) (Yuan et al., 2010) to attempt to identify the sequence type (ST) of the bacterium that was potentially transmitted to the plant. Every month, about five leaves, when they were available, were collected and tested by PCR (Minsavage et al., 1994) to check the evolution of the bacterium in the plants. The experiment was repeated in late June 2021. In total 250 *P. spumarius* were collected, mainly in Inca, on olive and almond tree plots. Five insects were netted per plant for a transmission period of five days (approx. 120 hours), except for five plants per species which received ten insects. After that, the same procedure of extraction and detection as in October 2020 was implemented. The experiments ended in October 2021, one year after the first transmission experiment and one *Xylella*-season after the second transmission test. The 20 cm twigs directly inoculated by the insect and the 20 cm ones above the inoculated sections were tested by real-time PCR of Harper et al. (2010) and then by nested MLST (Cesbron et al., 2020).

### 2.7 Statistical analysis

Statistical analyses were carried out to determine whether there were significant differences in bacterial population detected in the stem of the different plant species and between the temperature conditions. The statistics were made on the transformed data obtained by real-time PCR, expressed in potential log (CFU/g of plant tissue), using the open source software R© 4.0.4 (R Core Team, 2021). One-way analysis of variance (ANOVA) with significance at P < 0.05 was used to compare the population average detected in the entire inoculated branch (from the IP to the apex). A contrast matrix with the multcomp package (Hothorn et al., 2008) was built to undertake simultaneous tests on the different groups of interest (i.e. the different species in each temperature condition and the two temperature conditions within the same species). To compare bacterial population at the IP only as well as at the apex only, the data did not meet assumptions required by ANOVA and Kruskal-Wallis non-parametric test (P < 0.05) was used. Wilcoxon pairwise tests were then applied to compare the means of each specific group and p-values were corrected using the Benjamini-Hochberg procedure.

## 3 RESULTS

### 3.1 Detections through time

Detections were performed over 17 mpi on petioles and on branches. The number of positive plants detected through time can be visualized in Table 1 for each plant species and conditions. The bacterium was detected by PCR in 53 % of the plants at the petiole level already at 1 mpi (Table 1). At 9 mpi, the bacterium was detected in 100 % of the tested plants, either by real-time PCR or observed by confocal microscopy, except for *P. canescens* grown in the 22 °C greenhouse (detection in four out of seven plants – 57 %). Roots of one to four plants from each species and conditions also tested positive by quantitative PCR or confocal microscopy, except for *P. canescens* under 28 °C conditions. At 17 mpi, the bacterium was still detected in plants that had not been yet submitted to final destructive analysis, for all three species tested. It was also detected in a sucker (root sprout) of one *P. tremula* plant (Ct 26.31 with real-time PCR). The bacterium was successfully detected in the inoculated tobacco plants at 1 mpi and in the periwinkle plants at 1, 2 and 3 mpi, while consistent negative results were obtained when detection of the mock-inoculated and non-inoculated plants were performed.

**Table 1.** Number of positive plants on the total number of inoculated plants at different months after inoculation (mpi) of the *X. fastidiosa* KLN59.3 strain in four Salicaceae species

|  | T°<br>conditions | 1 mpi<br>Petiole<sup>a</sup> | 3 mpi<br>Petiole<sup>a</sup>/Branch<sup>b</sup> | 9 mpi<br>Branch<sup>c</sup> | Roots<sup>d</sup> | 17 mpi<br>Branch<sup>e</sup> |
| --- | --- | --- | --- | --- | --- | --- |
| *Populus tremula* | 22 °C | 5/10 | 8/10 | 6/6 (100 %) | 1/2 | 3/3 |
|  | 28 °C | 11/13 | 9/13<sup>b</sup> | 7/7 (100 %) | 3/3 | 0/1 |
| *Populus canescens* | 22 °C | 4/10 | 1/10 | 4/7 (57 %) | 1/1 | 1/3 |
|  | 28 °C | 2/10 | 7/10 | 7/7 (100 %) | 0/1 | 0/3 |
| *Salix alba* | 22 °C | 8/10 | 9/10 | 8/8 (100 %) | 2/2 | 2/2 |
|  | 28 °C | 4/13 | 12/13<sup>b</sup> | 7/7 (100 %) | 1/1 | 1/1 |
| *Salix caprea* | 22 °C | 3/6 | 0/6 | 6/6 (100 %) | 3/6 | - |
|  | 28 °C | 4/6 | 0/6 | 6/6 (100 %) | 4/6 | - |

<sup>a</sup> Group of plants for which bacterial detection was performed on petioles by PCR of Minsavage *et al.* (1994)
<sup>b</sup> Group of plants for which bacterial detection was performed for 10 plants on petioles by PCR of
Minsavage *et al.* (1994) and for three plants on branches by real-time PCR of Harper *et al.* (2010) <sup>c</sup> Group of plants for which bacterial detection was performed on branches by real-time PCR of Harper *et al.* (2010) and/or by confocal microscopy visualization of the GFP-marked strain in the stem vessel sections
<sup>d</sup> Group of plants for which bacterial detection was performed on roots by real-time PCR of Harper *et al.* (2010) and/or by confocal microscopy visualization of the GFP-marked strain in the root vessel sections
<sup>e</sup> Group of plants for which bacterial detection was performed on stems by real-time PCR of Harper *et al.* (2010)

### 3.2 Monitoring movement in the inoculated branch

The movement of the bacterium in the inoculated branch through the different months is represented in Fig.1. The dispersion of the bacterium from the inoculation point (IP) was expressed in cm (Fig.1a) or in percentage of the total branch length colonized by the bacterium (Fig.1b). The maximum dispersion recorded from the IP can also be observed in the same figure.

**Fig. 1.**
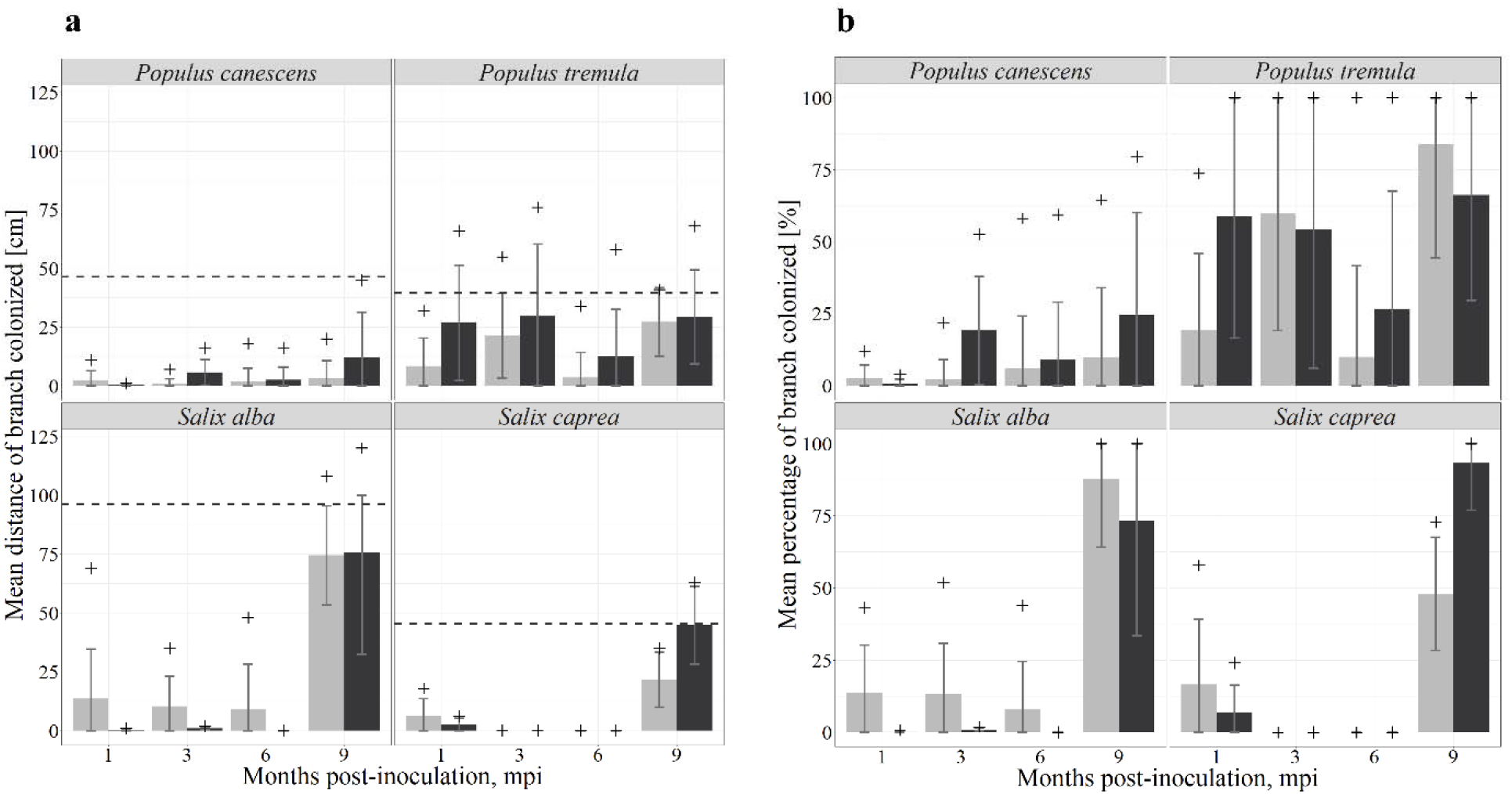
Evolution through the different months of the dispersion of the bacterium in the inoculated branches from the inoculation point for each species and each temperature condition, expressed in **a**. distance in cm and in **b**. percentage of branch colonized. The light grey bars represent the plant group placed at 22 °C and the dark grey ones the plant group placed at 28 °C. The “+” represents the maximal dispersion recorded for each group. The “-” on the ordinate-axis represent the average size of the inoculated branch. The dashed lines in **a**. represent the average total length of the inoculated branch at month 9 for each respective species, to put it in perspective with the length colonized by the bacterium. From month 1 to month 6, the dispersion was assessed through the bacterial detection in petioles by PCR (Minsavage *et al*., 1994), while at month 9 it was assessed through real-time PCR detection (Harper *et al*., 2010) in branches and confocal observation in stem sections.

*P. tremula* is the species that showed the most consistent detections away from the IP from month 1 to month 6. Already at 1 mpi, the bacterium was detected in petioles 65 cm away for two plants of the 28 °C-greenhouse, corresponding to a colonization of 100 % of the inoculated branch, as these petioles were located at the apex of the branches. The maximum distances were recorded at 3 mpi and were 55 cm at 22 °C and 76 cm at 28 °C. At 9 mpi, the bacterium had colonized 100 % of the inoculated branch for five plants out of seven at 22 °C and for three plants at 28 °C.

For the other species, the petiole detection technique showed less migration from month 1 to month 6. However, for *S. alba*, the bacterium was still detected in three individuals of the temperate conditions quite consistently across the different months, with already a detection in one individual 69 cm above IP after only 1 mpi. At 9 mpi, real-time detections in the stems highlighted a greater migration distance in *S. alba* and *S. caprea* than in the previous tested months. In *S. alba*, 100 % of the branch was colonized in six plants in the 22 °C-conditions and in four in the 28 °C-conditions, while there were respectively none and five for *S. caprea*. The greatest distances traveled were detected in *S. alba* with maximum distances above IP of 108 cm at 22 °C and 120 cm at 28 °C at 9 mpi. For *S. caprea*, the bacterium was detected until 35 cm above IP at 22 °C and 63 cm at 28 °C at 9 mpi. Less migration was measured in *P. canescens* for all mpi with still maximum distances of 20 cm at 22 °C and 45 cm at 28 °C at 9 mpi, but never reaching 100 % of branch colonization.

### 3.3 Quantification in plant tissues

The average of bacterial concentration at 9 mpi expressed in potential CFU/g of plant tissue in the different types of tissues for each species and each condition can be viewed in Table 2. The comparison of the log of these values is represented in boxplots in Fig. 2 for the entire inoculated branch (average of concentration detected from the IP to the apex), for the IP and for the apex portion of the branch. Fluorescence signal in the negative controls remained below the background, with no Ct values.

**Table 2.** Average of bacterial concentration 9 mpi expressed in potential CFU/g of plant tissue determined by real-time PCR (Harper et al., 2010) after the generation of a standard calibration curve, in different plant parts and for each species and each temperature growing condition. The apex, IP and average population in the entire inoculated branch (average of concentration detected from the inoculation point to the apex) were subjected to statistical analysis between both conditions and between the different species. Standard deviation was shown only for the average of population in the entire inoculated branch.

|  | *P. canescens* |  | *P. tremula* |  | *S. alba* |  | *S. caprea* |  |
| --- | --- | --- | --- | --- | --- | --- | --- | --- |
|  | 22 °C | 28 °C | 22 °C | 28 °C | 22 °C | 28 °C | 22 °C | 28 °C |
| Apex | 3.43E + 02 | 1.11E + 03 | 3.18E + 06 | 4.74E + 04 | 1.53E + 05 | 7.96E + 04 | 3.59E + 02 | 2.86E + 06 |
| Above IP | 6.88E + 02 | 4.11E + 03 | 8.44E + 06 | 9.16E + 04 | 1.05E + 05 | 2.37E + 05 | 1.59E + 05 | 3.61E + 06 |
| IP | 3.81E + 03 | 3.72E + 04 | 2.97E + 06 | 3.30E + 05 | 8.12E + 04 | 1.74E + 04 | 1.16E + 05 | 1.00E + 05 |
| Below IP | 4.09E + 04 | 2.77E + 03 | 1.67E + 06 | 2.12E + 05 | 1.49E + 04 | 3.86E + 03 | - | - |
| Roots | 2.77E + 05 | 0.00E + 00 | 4.38E + 05 | 1.52E + 06 | 1.05E + 05 | 3.92E + 02 | 1.80E + 05 | 4.35E + 04 |
| Total in inoculated branch | 1.52E + 03 ± 2.08E + 03 | 1.28E + 04 ± 1.10E + 04 | 3.56E + 06 ± 5.64E + 06 | 2.63E + 05 ± 3.76E + 05 | 1.19E + 05 ± 6.80E + 04 | 1.43E + 05 ± 3.41E + 05 | 9.32E + 04 ± 1.07E + 05 | 2.08E + 06 ± 1.70E + 06 |

**Fig. 2.**
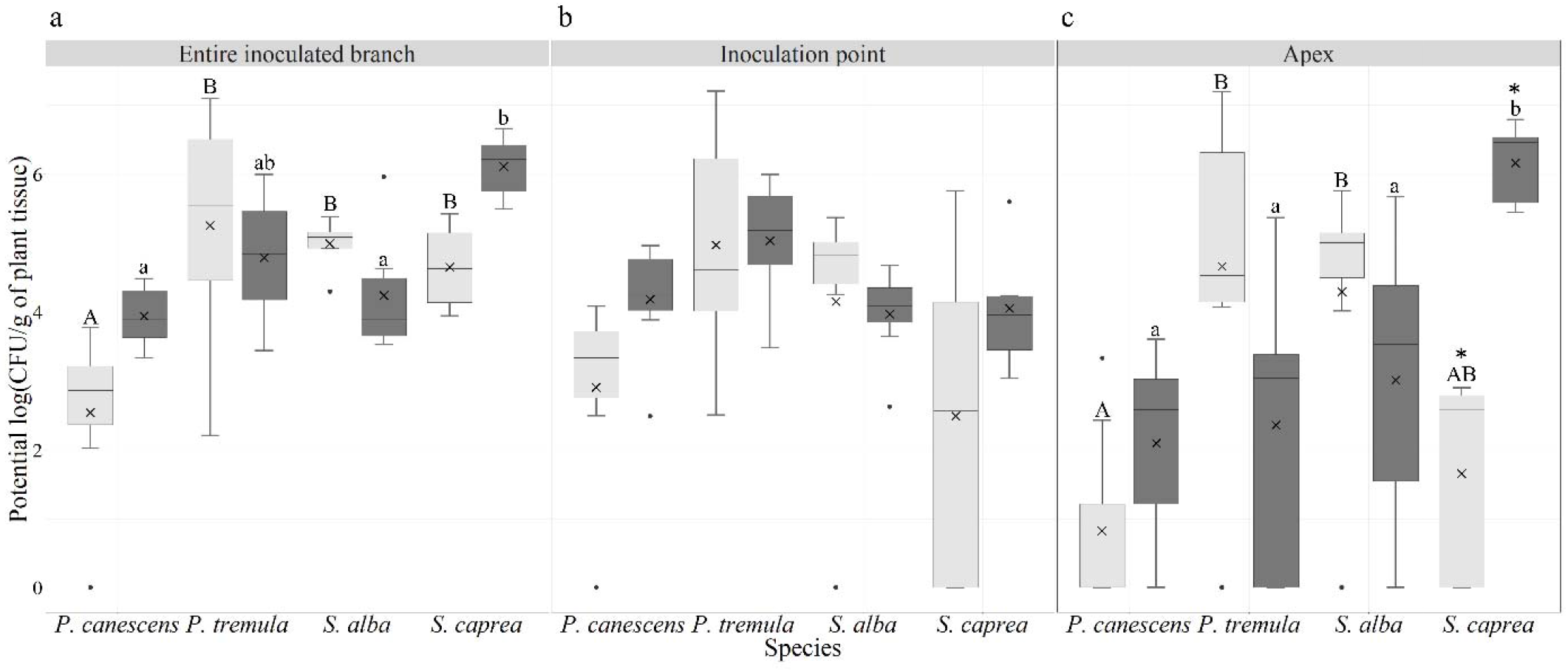
Comparison of the average of bacterial concentration 9 mpi, expressed in potential log (CFU/g of plant tissue) determined by real-time PCR (Harper et al., 2010) after the generation of a standard calibration curve, for each species and each temperature condition in different plant parts: **a**. in the entire inoculated branch (average of concentration detected from the inoculation point to the apex), **b**. at the inoculation point and **c**. at the apex of the branch. The light-grey boxes represent the data from the plants maintained at a temperature of 22 °C and the dark-grey boxes from the plants maintained at 28 °C. The horizontal black lines denote median values, boxes extend from the 25th to the 75th percentile of each group’s distribution of values, and vertical extending lines denote the range of values, “x” represent the mean for each group. The capital letters refer to significant differences between the four species in the 22 °C-greenhouse, and the lowercase letters refer to significant differences between species in the 28 °C-greenhouse. Groups accompanied by the same letter are not significantly different. * indicates significant differences between the two conditions in the same species. Statistical analyses were done by ANOVA and Tukey means comparison tests in graph **a**. and by Kruskal-Wallis followed by Wilcoxon test for paired data in graphs **b**. and **c**

When comparing both temperature conditions for each species, pairwise tests only indicated a significant difference for *S. caprea* at the apex (p = 0.038), the detected bacterial population being higher at 28 °C than at 22 °C. For *P. canescens*, the same pattern seems to occur with higher bacterial populations at 28 °C. In contrast, for *S. alba* and *P. tremula* the opposite pattern was observed with higher concentrations at 22 °C; however, these differences were not statistically confirmed.

A comparison was made between the four species under the same temperature growing conditions. In the entire inoculated branch, significant differences were measured between the four species (p < 0.0001). At 22°C, pairwise tests indicated that *P. canescens* had lower concentrations in their entire branches than *P. tremula* (p < 0.001), *S. alba* (p < 0.001) and *S. caprea* (p < 0.001); at 28°C pairwise tests measured that *S. caprea* had significantly higher concentrations than *S. alba* (p = 0.013) and *P. canescens* (p = 0.003). These differences were mainly due to different amounts detected at the apex (p = 0.0005). There, pairwise tests showed that concentrations were significantly lower for *P. canescens* than for *P. tremula* (p = 0.038) and *S. alba* (p = 0.038) at 22°C; while at 28°C, *S. caprea* had significantly higher concentrations at the apex than *S. alba* (p = 0.039), *P. tremula* (p = 0.038) and *P. canescens* (p = 0.038). No significant differences were measured between the four species at IP despite high values detected in *P. tremula* reaching about 1.6E + 07 CFU/g of plant tissue in one plant at 22 °C.

Bacteria were also detected below IP, as well as in the roots with the greatest quantities in *P. tremula* at 22 °C reaching an average of 1E + 06 CFU/g (Table 2). Bacteria in roots were detected for all groups, except for *P. canescens* at 28 °C. However, in this last group, bacteria were still detected 20 cm below the IP in the stem.

### 3.4 Multiplication of bacteria

Quantification of *X. fastidiosa* allowed to compare the potential number of total CFU detected in one plant with the number of CFU initially inoculated in the branch (max 5.5E + 06 CFU or 1.1E + 07 CFU if two branches were inoculated) to check if bacterial multiplication in the plant occurred. Such multiplication has been measured with certainty in three *P. tremula* at 22°C, one *S. alba* at 28 °C and four *S. caprea* at 28 °C.

### 3.5 Bacteria distribution in xylem vessels

In the periwinkles, bacteria were observed in a series of adjacent vessels (Fig. 3), either in clusters completely or partially obstructing these vessels, or in single-cells free in the lumen or attached to the vessel walls.

**Fig. 3.**
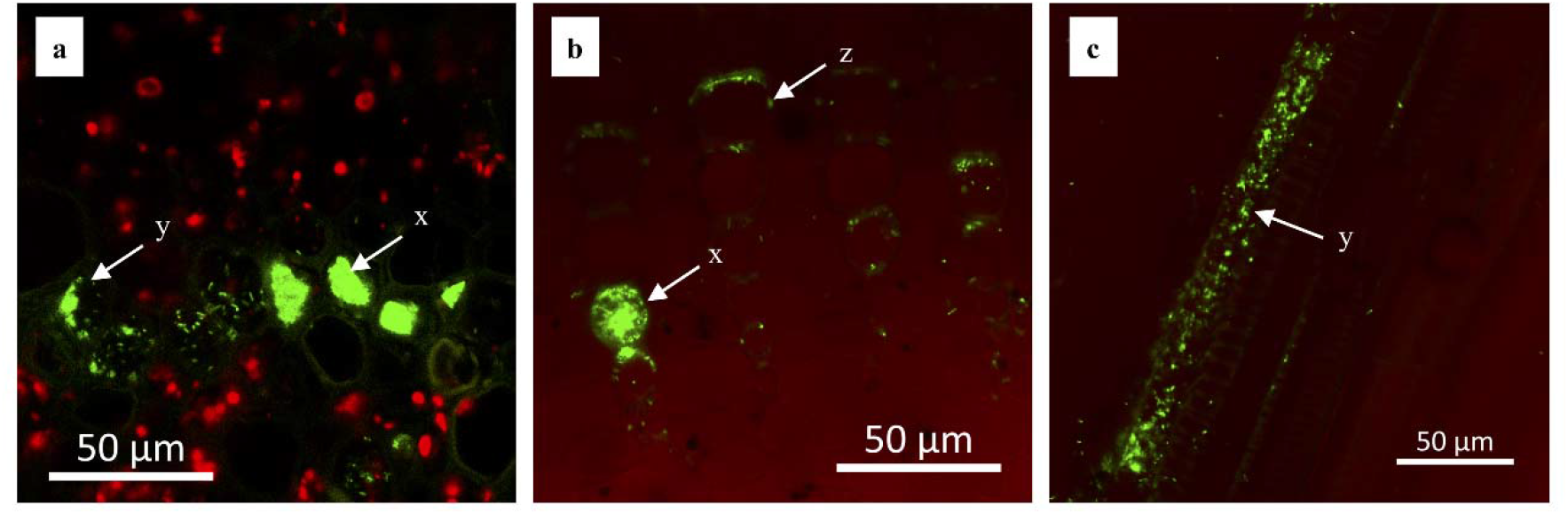
Transversal sections (a,b) and longitudinal (c) sections of *Catharanthus roseus* stems at the inoculation point (IP), inoculated with *X. fastidiosa* KLN59.3 expressing GFP observed by confocal microscopy. *X. fastidiosa* cells contained in xylem vessels are depicted in green; plant tissues and xylem are depicted in red. Bars represent a 50 µm scale. x: obstructed vessels, y: cells aggregates partially obstructing vessels ; z: single cells attached to the vessel wall.

In *P. tremula, X. fastidiosa* was observed in all inoculated plants, at every IP and at the apexes of 5/8 plants (62.5 %). Bacteria were observed in different forms (Fig. 4): single-cells in the lumen or clinging to the vessel walls, or in clusters attached to the wall, partially or totally obstructing the vessels. Generally, a combination of these different forms could be observed in all the sections. Close to the IP, many clogged vessels were pointed out. Moving upwards from the IP, two different situations were observed. The first was a reduction in the number of obstructed vessels, and a decrease in the number of bacteria as one approached the apex. For three *P. tremula* plants, the opposite situation was observed, with an increase in the number of blocked vessels the closer one got to the tip of the stem, most vessels being totally blocked on the transversal sections at the apex (Fig. 4f). The bacteria were also consistently observed in the roots and in the stem below the IP of *P. tremula*, where a high proportion of vessels were filled with single-cells or partially obstructed by bacterial aggregates. Nonetheless, in two plants at 28°C, a high proportion of totally obstructed vessels were observed as well. An overview of the distribution of *X. fastidiosa* in all the plant parts observed in the transversal sections is schematized in the Supplementary Fig. S1. These diagrams allow highlighting patterns of bacterial colonization in the plant. No specific patterns were highlighted for *P. tremula*, the colonized vessels being distributed randomly in the sections. In general, there were large differences between individuals grown under the same conditions, ranging from low to high colonization. On these diagrams, more totally and partially obstructed vessels were pointed in the inoculated stems of *P. tremula* maintained at 22°C, while they were greater in the root sections of the plants grown at 28°C.

**Fig. 4.**
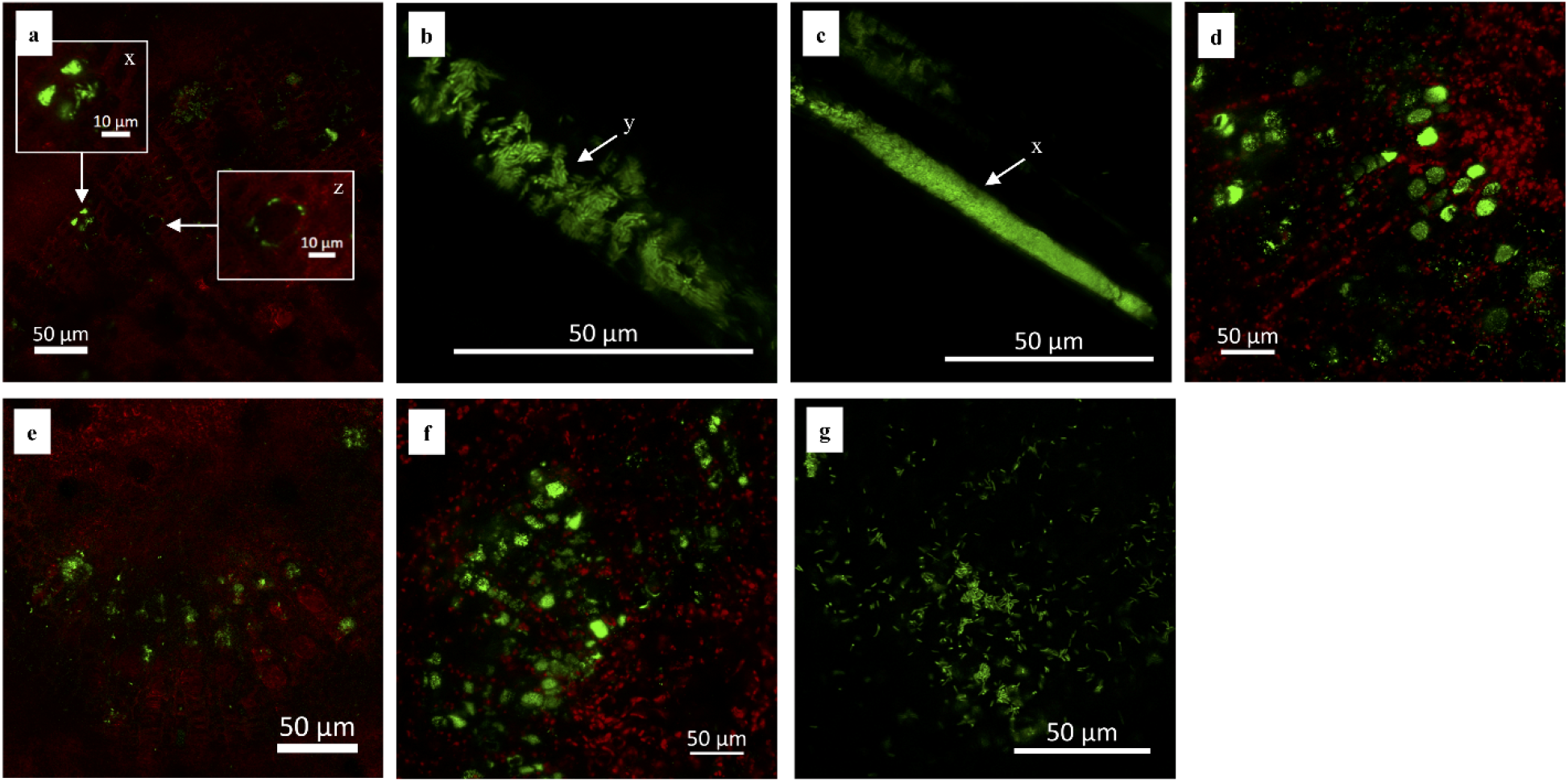
Transversal sections and longitudinal sections of *Populus tremula* stems inoculated with *X. fastidiosa* KLN59.3 expressing GFP observed by confocal microscopy. *X. fastidiosa* cells contained in xylem vessels are depicted in green; plant tissues and xylem are depicted in red. Bars represent a 50 µm scale, except in the zoom in a) where it represents a 10 µm scale. x: obstructed vessels, y: cells aggregates partially obstructing vessels; z: single cells attached to the vessel wall. a) transversal-section in the stem at the inoculation point (IP); b) & c) longitudinal-section in the stem at the IP; d) transversal-section in the stem 9 cm above the IP; e) transversal-section in the stem at the apex, 26 cm from IP; f) transversal-section in the stem at the apex, 12 cm from IP; g) transversal-section in the roots, 48 cm below IP.

Aggregates and biofilms of *X. fastidiosa* could also be observed by scanning electron microscopy in transversal sections of *P. tremula* (Supplementary Fig. S2), being the only species for which *X. fastidiosa* could be spotted by this method.

For *S. alba* in both temperature regimes, the fluorescent bacteria (Fig. 5) were often visible as single-cells isolated in the lumen of the vessels or attached to the walls, sometimes in small clusters, but very rarely blocking the vessels. The bacteria were also spotted in less xylem vessels than for *P. tremula*. It was detected at IP of all the plants observed at 22°C and of 1/3 plants observed at 28°C. However, for the remaining two, the bacterium was observed higher in the stem, at 54 cm and 75 cm from the IP. It was never spotted at the apex, however, since the stems of *S. alba* were long, the bacterium was still observed at a maximal distance of 92 cm from the IP at 22°C and 81 cm at 28°C. Similarly, *X. fastidiosa* also moved below the IP, observed up to 19 cm-below at 22°C and 26 cm at 28°C, and in the roots of one plant at 28°C. In the Supplementary Fig. S1, partially obstructed vessels were also more numerously observed in the sections of the inoculated branches maintained at 22°C, while no differences could be made below the IP and in the roots. The colonized vessels were either isolated or adjacent to other colonized vessels. In two plants, they were located only in one side of the section.

**Fig. 5.**
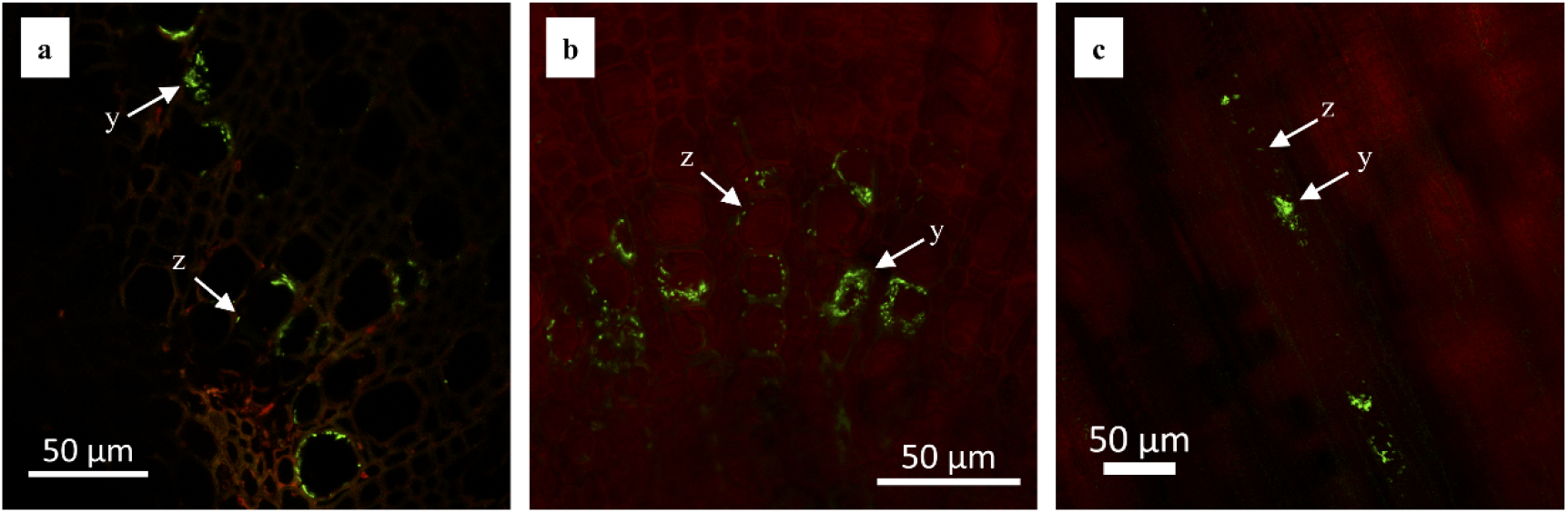
Transversal sections and longitudinal sections of *Salix alba* stems inoculated with *X. fastidiosa* KLN59.3 expressing GFP observed by confocal microscopy. *X. fastidiosa* cells contained in xylem vessels are depicted in green; plant tissues and xylem are depicted in red. Bars represent a 50 µm scale. y: cells aggregate partially obstructing vessels; z: single cells attached to the vessel wall. a) transversal-section in the stem 5 cm above the inoculation point (IP); b) transversal-section in the stem 42 cm above the IP; c) longitudinal-section in the stem 25 cm above the IP.

In *P. canescens, X. fastidiosa* was visible only at the IP, in 1/3 plants observed at 22°C and 2/3 observed at 28°C, but in very small amounts, and never completely clogging the vessels.

Due to the delay in inoculation of *S. caprea*, confocal microscopy could not be performed on this species.

### 3.6 Bacterial isolation

The bacterium could be isolated on BCYE medium from *P. tremula*, from four plants maintained at 28 °C at 4 mpi and 9 mpi (from IP and apex) and from one plant maintained at 22 °C at 9 mpi (from IP and apex). For *S. alba*, it was successfully isolated from one plant at 28 °C at 4 mpi (only at the IP). Colonies grew slowly, requiring at least 15 days to become visible and all of them were confirmed as *X. fastidiosa* by PCR and Sanger sequencing. Isolation was also obtained for every inoculated periwinkle. No colonies could be reisolated from *P. canescens* out of 14 attempts.

### 3.7 Symptom monitoring

At 1 mpi, at both 22 °C and 28 °C conditions, light chlorosis and/or necrosis developed, either at the leaf margins for *P. tremula*, or on the entire leaf surface for the other species. Nevertheless, such symptoms were also detected, although to a lesser extent, on the uninoculated control plants, probably because our experimental setup was constrained by the biosafety containment measures. Seven months after inoculation, a generalized high defoliation started to occur on all plants, controls included.

For *P. tremula*, clear symptoms of desiccation could be highlighted for the inoculated group in both conditions. The plant desiccation most often started from the apical portion of the inoculated shoots and then could diffuse to the entire plant. The symptoms recorded were ranked into five categories according to their severity (Fig. 6a). For each plant, the evolution of these symptoms at 1, 4, 7 mpi has been represented in Fig. 6b, where it can be observed that a minority of plants have already shown severe symptoms at their inoculated branch at 1 mpi. At 4 mpi, half of the plants had heavily affected inoculated branches at both 22 °C and 28 °C. At 7 mpi, more than half of the inoculated branches were completely dead at 28 °C but most of the plants had only mild symptoms on the rest of their foliage. At 22 °C, most of the plants had all the leaves of their inoculated branches completely necrotic, but the wood of the stem was still alive. However, in that greenhouse, the rest of the foliage was more affected compared to the individuals at 28 °C. The latency period in *P. tremula* for both conditions was comprised between 1 and 4 months.

**Fig. 6.**
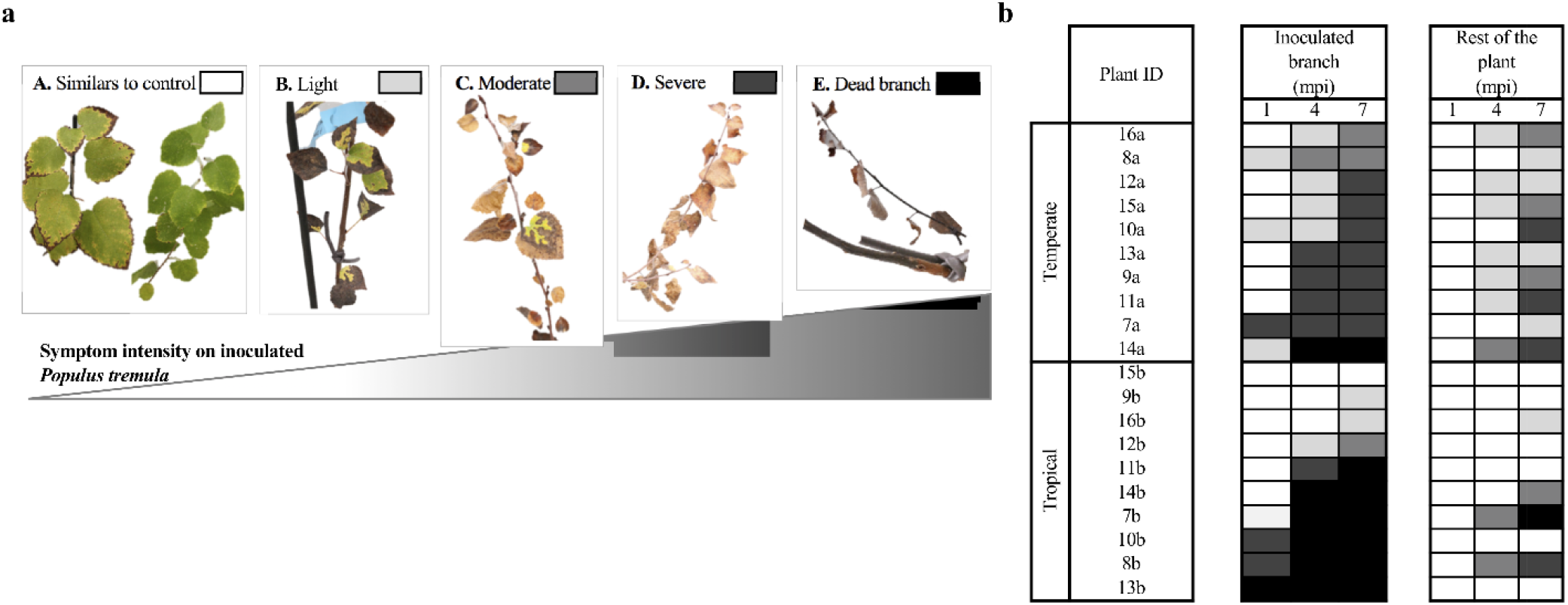
Symptoms on *P. tremula* after pin-pricked inoculation with *X. fastidiosa* KLN 59.3 strain. **a**. Categories of symptom severity ranked by intensity: A. symptoms similar to control group (uninoculated or mock-inoculated); B. light symptoms compared to the control group, with necrosis of leaf margins and less than half of the leaves completely necrotic attached to the branch; C. moderate symptoms with necrosis of leaf margins and more than half of the leaves completely necrotic attached to the branch; D. severe symptoms with all the leaves of the branch completely necrotic but with a still-living stem; E. Completely dead branch. **b**. The evolution of the symptoms 1, 4 and 7 mpi on each inoculated individual from temperate (22 °C) and tropical (28 °C) greenhouse, for their inoculated branch and for the foliage and stem on the rest of the plant. This evolution is represented thanks to the legend in Fig. 4a. The individuals are classified in each greenhouse from the less to the most symptomatic inoculated branch.

Symptoms different from the control group also appeared on eight inoculated *S. caprea*. At 2 mpi, four plants at 28 °C showed several successive dried leaves starting from the IP. The symptoms recorded at 8 mpi were ranked according to their severity and are illustrated in Fig. 7a. At 8 mpi, two plants at 22 °C and three at 28 °C had one of their inoculated branches completely desiccated and dead (Fig. 7b). Furthermore, three plants at 28 °C and two at 22 °C had at least 50 % of their total inoculated branch length defoliated starting from IP.

**Fig. 7.**
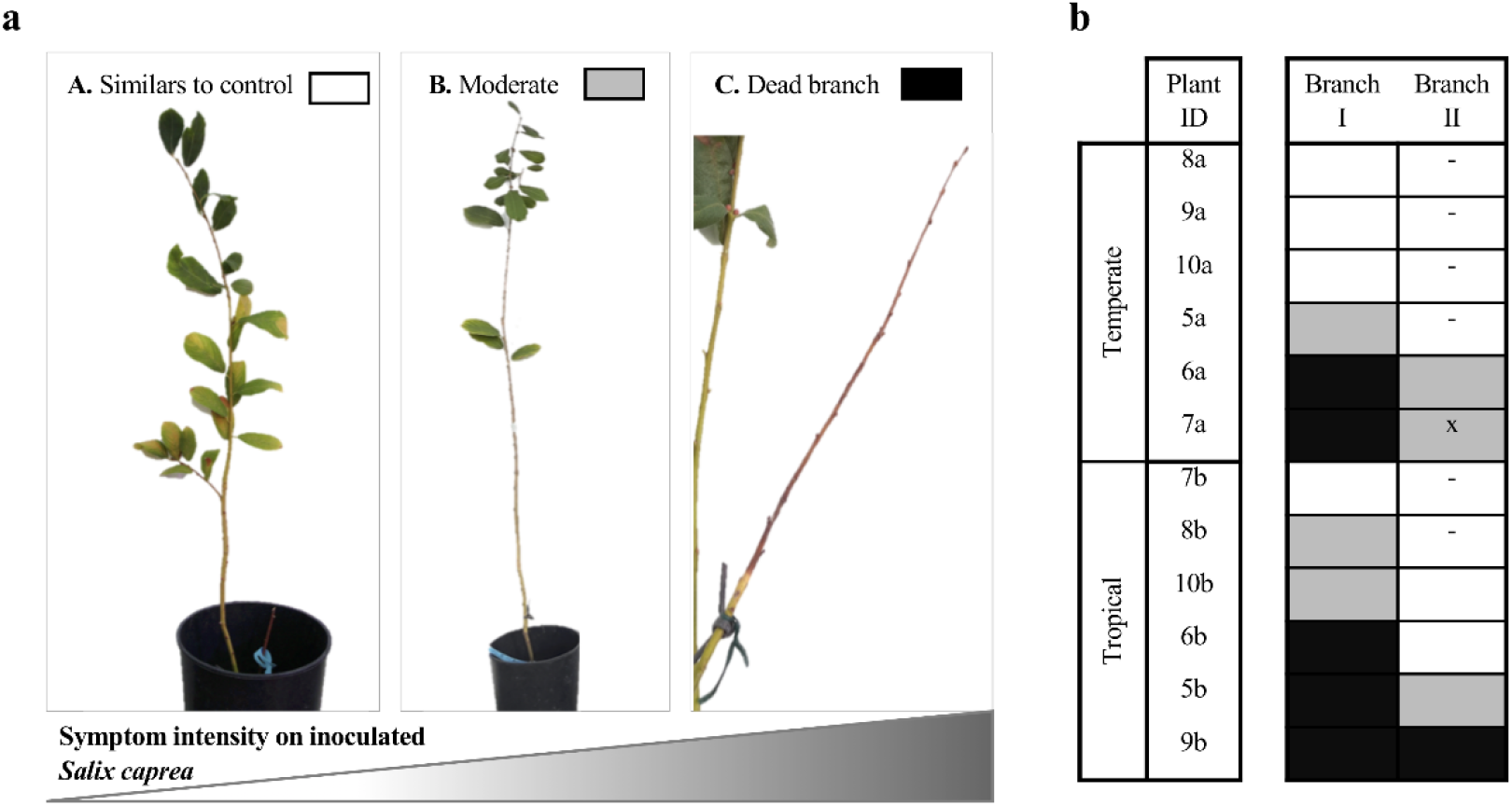
Symptoms on *S. caprea* after pin-pricked inoculation with *X. fastidiosa* KLN 59.3 strain 8 mpi. **a**. Categories of symptom severity ranked by intensity: A. symptoms similar to control group (uninoculated or mock-inoculated); B. moderate symptoms compared to the control group, defoliation of at least 50 % of the total inoculated branch length starting from the inoculation point; C. Completely dead branch. **b**. The symptoms 8 mpi on each inoculated individual from temperate (22°C) and tropical (28°C) greenhouse, for their two inoculated branches “Branch I” and “Branch II”. The intensity of symptoms for each individual is represented thanks to the legend in Fig. 5a. “-” in the symptom intensity box means that there were no “Branch II” (only one inoculated branch) ; “x” means that this second branch was not an inoculated one. The individuals are classified in each greenhouse from the less to the most symptomatic.

From the inoculation date to the final destruction of the inoculated branches, no typical symptoms could be highlighted for *P. canescens* and *S. alba* in both temperature conditions when compared to the control group. For all species, no significant differences in terms of growth were measured between the various controls and inoculated groups

### 3.8 Transmission experiments in Mallorca

During the transmission period, *P. spumarius* were observed feeding on the plants, as feeding fluid was observed in most of the nets. The mortality rate of the insects at the end of the transmission period in October 2021 was 11.5 %. Out of the 200 *P. spumarius*, 32 had a Ct < 35 and were considered positive, leading to a prevalence rate of 16 %. Of these 32 insects, 28 had a Ct between 25 and 28. There were undetermined results for five insects, with a Ct between 35 and 38. The ST of 33/37 positive or undetermined insects could be identified, being ST81 (*X. fastidiosa* subsp. *multiplex*) for 23 insects (70 %) and ST1 (*X. fastidiosa* subsp. *fastidiosa*) for 10 insects (30 %). One year after the transmission test, the bacterium was detected in a single *S. alba* plant by real-time PCR (Ct 24.57 ± 0.08) and identified as ST81, corresponding to the ST identified in the respective insects. In this plant, the bacterium was identified in the inoculated branch section and in the upper section, 20 cm above the inoculated portion. In total, insects were found positive on 10 *S. alba* plants, leading to 1/6 infected plant among ST81 inoculated ones, and 0/4 infected plant among ST1 inoculated ones.

In the June 2021 transmission experiment, the observed insect mortality was 7 %. Only 13 out of 250 insects were detected positive for the bacteria (5.2 %). In addition, MLST performed successfully on nine samples, allowed the exclusive identification of the sequence type ST1. Four months after the transmission, ST1 bacteria were detected in two *S. alba* plants, at the inoculated section but also in an upper section, 20 cm above (Ct 24.14 ± 0.15 and 27.26 ± 0.51). In total, insects were found positive on five *S. alba*, leading to a ratio of two infected plants out of a total number of five inoculated ones.

All the potentially inoculated branches showed a high defoliation. However, this defoliation was observed on most of the plants and therefore, could not be definitely linked to the presence of the bacterium. None of the *P. tremula* plants were detected positive after both transmission experiments, with ratios of 0/4 plants for ST81 and of 0/4 for ST1 in October, and of 0/3 for ST1 in June.

## 4 DISCUSSION

### 4.1 Need to evaluate the potential new host plants for *X. fastidiosa* in the EU

*X. fastidiosa* is probably the plant pathogenic bacterium with the widest host range reported to date, with up to 655 host species reported so far (EFSA, 2022). The recent discovery of *X. fastidiosa* in Europe raised the question of the susceptibility of indigenous European host plant species, as well as the bacterium ability to spread further North in Europe and to establish unforeseen tripartite interaction between both European insect vectors and host species (EFSA, 2019).

Here a focus was given on potential riparian host plants, based on the role played by such habitat for the maintenance of *X. fastidiosa* infested plants in the environment and common presence of potential vectors (Freitag, 1951; Purcell, 1974). Salicaceae species have been investigated in California, as perennial host of riparian areas (Purcell and Saunders, 1999). In this study, we showed that Salicaceae species like *Populus tremula*, the Eurasian aspen, or *Salix alba*, the white willow, despite their Eurasian origin and distribution, and although they have not been exposed to *X. fastidiosa* in the past, are potential hosts for the bacteria.

### 4.2 *Populus tremula* as a potential host plants for *X. fastidiosa*

The susceptibility of *P. tremula* has been confirmed by mechanical inoculation of the GFP-tagged *X. fastidiosa* subsp. *fastidiosa* Temecula strain, followed by confirmation of the distribution of the bacteria in the xylem vessels of inoculated plants along the branches, over a period up to 17 months. Bacteria have been evidenced by PCR, real-time quantitative PCR, by confocal and scanning electron microscopy. *Xylella fastidiosa* has been re-isolated successfully several times from inoculated *P. tremula* plants at 4 and 9 mpi. Purcell and Saunders (1999) have isolated *X. fastidiosa* from another *Populus* species, Fremont cottonwood (*Populus fremontii*) 30 days following insect-vector inoculations. They also reported similar ranges of CFU per gram of plants as those recorded in this study (i.e. log 4-6).

Despite the biosafety conditions inherent to the quarantine status of *X. fastidiosa*, symptoms have been recorded, but should be regarded with caution since they were observed under greenhouse growing conditions. *Populus tremula* was the most susceptible species, with more than half of the individuals showing advanced symptoms of desiccation, starting most often from the tip of the inoculated branches and strongly linked with *X. fastidiosa* inoculation. This is also related with the highest levels of CFU recorded per inoculated plants and to the number of bacteria-obstructed xylem vessels in the stem above the inoculation point. The species developing symptoms are often those with the largest bacterial populations at the inoculation point, in line with studies carried out with other plant species by several authors (Hill and Purcell, 1995; Holland et al., 2014; Saponari et al., 2017). In their study, Saponari et al. (2017) showed in olive that the cultivar developing the highest symptomatic response was the one with the largest bacterial populations with a range of CFU/ml detected by real-time PCR similar to the one observed here.

### 4.3 *Salix alba* is also a host for *X. fastidiosa*, but without the evidence of symptom production

In our hands, *S. alba* is also a host for *X. fastidiosa*, yet the bacterial population and circulation in *S. alba* is not leading to diseased plants. Some *X. fastidiosa* hosts may show moderate symptoms when grown in the greenhouse, but none when grown in the field (Purcell, 2013). The ability to circulate significantly but at low density and to distribute randomly in xylem vessels, thanks to an enzymatic degradation of pit-membrane (Roper et al., 2007) probably allows the bacterium to avoid harming its host and to establish an endophytic interaction with *S. alba*, as it occurs with other plant species (Chatterjee et al., 2008).

Purcell and Saunders (1999) have proposed that the infection of *X. fastidiosa* in red or arroyo willow is transient, with a multiplication in the first weeks following infection, followed by a diminution after 12 weeks and a possible death of the bacteria. Here the bacteria were systematically detected in *S. alba* after nine months, and were also detected in the single plant tested at 17 mpi. Interestingly, we have been able to isolate *X. fastidiosa* from *S. alba* after 4 mpi, but not after nine months. It is, however, complicate to ensure that the randomly distributed parts colonized by the bacteria have been sampled.

### 4.4 Susceptibility to *X. fastidiosa* is different among Salicaceae – *P. canescens* and *S. caprea*

*P. canescens* and *S. caprea* have also been investigated as potential host plants. Here again, we have been able to detect the bacteria in both plant species by quantitative real time PCR and confocal microscopy.

*Salix caprea* behave very similarly to *S. alba* with comparable load detected at 22°C. However, the bacterium seems to multiply more in *S. caprea* placed at 28°C with higher load and with more symptoms observed.

On the other hand, the number of CFU detected in *P. canescens* was the lowest (log 3-4), with numbers probably below the threshold needed for a possible transmission by insect vectors, which was measured at log 4 CFU/g of plant tissue for grape by Hill and Purcell (1997). This highlights once again the differences in susceptibility between plants of different varieties or closely related species. Several authors had already reported significantly different levels of infection within the same genus, e.g. in *Prunus* or *Vitis* (Hao et al., 2016; Ledbetter and Lee 2018), or between different cultivars of the same species, as in olive or blueberry (Saponari et al., 2017; Burbank et al., 2020). What will trigger symptoms in one species or cultivar and not in another remains poorly understood and varies according to the host-strain combination (EFSA, 2015; Nunney et al., 2019). According to studies on coffee, plum and grapevine, the severity of symptoms is associated with the proportion of bacterial colonization of xylem vessels (Newman et al., 2003; Alves et al., 2004; Baccari and Lindow 2011). Here, for *P. canescens*, blocked vessels have never been seen in confocal microscopy.

### 4.5 Insect transmission and susceptibility to different strains of *X. fastidiosa* subspecies

The susceptibility of plants also varies according to the bacterial strain (Nunney et al., 2019) and carrying out trials with other strains would certainly be useful. In a preliminary test, a different strain from the subspecies *fastidiosa* (LMG17159) was inoculated in *P. tremula* and in *S. alba*, with positive detection later on (data not shown). Here the focus was given to the subspecies *fastidiosa* and the Temecula strain, considering the advantage of assessing host plant susceptibility with the support of confocal microscopy. Investigations on some *multiplex* subsp. strains are also necessary, as the subspecies *multiplex* was found in northern regions of America and could therefore be more adapted to northern temperate European areas (EFSA, 2019). The Mallorcan transmission experiments using field collected *P. spumarius*, besides investigating the major role of insect vectors, also allowed to test two other strains on *P. tremula* and *S. alba*, another ST1 subspecies *fastidiosa* and ST81 subspecies *multiplex*. Here, the transmission by *P. spumarius* on *S. alba* has been proven possible, with 1/6 plants infected by *X. fastidiosa* subsp. *multiplex* ST81 and 2/9 plants infected by *X. fastidiosa* subsp. *fastidiosa* ST1.

Although transmission tests with insects are much closer to natural conditions than mechanical inoculations, the length of the inoculation period affects its efficiency (Purcell and Finlay, 1979). By extending the inoculation period, the cells attached to the insect mouthparts would be more likely to detach and be inoculated in the plant (Almeida et al., 2005). Under natural conditions, insects have access to the plant throughout all the season while only for a period of 4-5 days in this study. Even if fewer than 200 viable bacterial cells per insect would be enough for successful transmission of *X. fastidiosa* (Redak et al., 2004), the introduced concentration could have been too low in this trial to be detectable just after one season of bacterial multiplication. As for the October 2020 transmission, the plants went into dormancy immediately afterwards and therefore, the period was less favorable for the subsequent bacterial development in the plants. However, October is a suitable month to collect infected insect vectors in Mallorca as it was shown that in general, prevalence was lower in summer months (June-July) compared to autumn ones (October), with an average of 23% (López-Mercadal et al., 2021). The *X. fastidiosa* prevalence found in field-collected insects in October 2020 was higher (16%) than the one found in June 2021 (5.2%) in accordance with the values commonly found in the area. The prevalence of *X. fastidiosa* in insects may also favor bacterial transmission to plants. For example, Cornara et al. (2017) reported that in Italy the increasing of the prevalence found in *P. spumarius* in late summer/ early autumn months, easily reaching 90%, enhances the potential transmission to plants.

In natural conditions, *X. fastidiosa* infection depends therefore on the frequency and abundance of insect vector feeding on a host. Salicaceae are host plants of several potential insect vectors in Belgium, including the polyphagous species *P. spumarius* or *A. alni*, which feeds on the four selected species (Hasbroucq et al., 2020). In addition, *A. salicina* can be extremely abundant on *S. alba* and on *S. caprea* in riparian areas (unpublished data). The transmission efficiency of these potential insect vectors still must be investigated. While some authors report that in order to be acquired, the bacteria should be in an “adhesive” state and thus rather in the form of a biofilm (Chatterjee et al., 2008), others state that biofilm would not be necessary for this acquisition (Almeida et al., 2005). Furthermore, the systemic movement of the bacterium in the xylem vessels would increases the ability of plants to serve as inoculum sources, independently of the vector activity (Purcell and Saunders, 1999).

### 4.6 Temperature effect and presence in roots and sprouts

While higher bacterial load were detected at 28°C in *P. canescens* and *S. caprea*, interestingly, for *P. tremula* and *S. alba*, the quantity of bacteria evidenced at 22 °C was higher than at 28 °C. Feil and Purcell (2001) have investigated the effect of a range of different temperatures on the *X. fastidiosa* growth in vitro, showing a higher multiplication at 28 °C compared to lower temperature. However, the authors stress also the fact that many factors influence the development of the bacteria in planta like plant growth stimulants or inhibitors (Davis 1978; Hopkins 1985). On the other hand, Roman-Ecija et al. (2021) reported relatively similar growth and biofilm formation in vitro of bacteria incubated at temperatures ranging from 16 to 32°C. If confirmed, the better development of *X. fastidiosa* at a lower temperature (22 °C compared to 28 °C) also supports the possibility of a potential role of *P. tremula* and *S. alba* further North in cooler European areas.

Furthermore, upward and downward migration to the root system has been observed for all species with higher concentrations for *P. tremula*. Roots would be less exposed to cold temperatures (Roman-Ecija et al., 2021) and could be a shelter for the bacterium to survive in colder regions, avoiding a potential winter curing. Bacteria were already reported in the root system including peach, blueberry and olive trees (Aldrich et al., 1992; Holland et al., 2014; Saponari et al., 2017), where higher bacterial populations were found in the most susceptible cultivars. The detection of the bacterium in one *P. tremula* sucker indicates that *X. fastidiosa* could be transmitted to new emerging seedlings through roots. Transmission by root grafting had already been reported on sweet orange (Hutchins et al., 1953; He et al., 2007). This could further increase the risk of dissemination through these pioneer species whose vegetative multiplication through sprouts is common in natural reservoirs, which could generate a pool of new infected seedlings at the end of winter.

### 4.7 Frequent hybridization of willows and poplars creating new genotypes

Hybridization being very common within the Salicaceae under natural conditions, the susceptibility variation in this hybrid population must be considered from an epidemiological point of view, either mitigating or intensifying the bacterial reservoir in natural or semi-natural areas. Several studies have investigated the susceptibility to different pathogens between the parental species and their natural or artificial hybrids. The response is influenced by environmental factors and varies according to the pathogenic agent (Fritz et al. 1999). Hybrid susceptibility can either be: i) similar to parents, or as one parent if they have different resistance traits and dominance occurs; ii) additive with an intermediate behavior between both parents; iii) lower if heterosis phenomena occur; iv) higher for example if the expression of resistance mechanisms is disrupted by recombination (Orians & Floyd, 1997; Fritz et al., 1999). After gathering information from 28 studies Fritz et al. (1999) reported the dominance of susceptible parents and susceptibility as the most common patterns.

*P. canescens* being a hybrid between *P. tremula* x *Populus alba*, it seems that hybridization has brought resistance traits to the Temecula strain in this species. Whether this difference in susceptibility comes from resistance traits in *P. alba* is not known. Interestingly, Pinon & Valadon (1997) reported *P. tremula* as a susceptible host of the bacterial canker caused by *Xanthomonas populi* (Xanthomonadaceae), bacteria from the same family as *X. fastidiosa*, while they indicate the hybrid between *P. tremula* x *P. alba* more tolerant and the parent species *P. alba* resistant.

## 5 CONCLUSION

This study highlighted two new potential hosts for the northern temperate European areas: *P. tremula*, highly symptomatic and *S. alba* rather asymptomatic. Asymptomatic plants make surveillance and early detection of the bacteria difficult, as visual inspection is not sufficient, and these plants must not be underestimated as they may spread the bacterium silently. *Salix caprea* seems to be a suitable host for the strain KLN59.3, however, more testing and isolation should be attempted to confirm this result. Finally, *P. canescens* seems, on the other hand, to be a species quite resistant to the inoculated strain. These results demonstrate once again that the *X. fastidiosa* host plant status cannot be predicted from the host family or genus, highlighting the importance of investigations on potential new hosts in areas where the bacterium is not known to occur. It also shows that, besides nurseries, natural environments such as riparian areas, should also be inspected for surveillance and early detection of new emerging diseases.

## ACKNOWLEGEMENT

We would like to acknowledge the SPF Santé publique for the long-term financial support through the two two-year-long projects: XYLERIS and XFAST. We are grateful to Pr. Steven E. Lindow (UC Berkeley) for kindly providing us with the KLN59.3 GFP-labelled strain. We would like also to thank Marie-Christine Eloy and Delphine Magnin (UCLouvain) for the technical support in confocal and scanning electron microscopy, Brigitte Vanpee for the plant care in the UCLouvain greenhouses and the UIB ZAP group, Pau Mercadal, Aroa Rodríguez and Carlos Barceló, for the help during transmission experiments including plant care and insect collects.

## AUTHOR CONTRIBUTIONS

LP, AG, AE, SH and NC contributed to the mechanical inoculation studies. JLM, MAM, SH and NC contributed to the insect transmission tests. CB, JCG and NC designed the study. The first and last authors wrote the manuscript. All coauthors commented on previous versions of the manuscript. All authors read and approved the final manuscript.

## FUNDING

The research that yielded these results, was funded by the Belgian Federal Public Service of Health, Food Chain Safety and Environment through the contracts RF 19/6331 (XFAST project) and RT/7 XYLERIS 1 (XYLERIS project). The research was also supported by the National Fund for Scientific Research (FNRS) and the Université catholique de Louvain (UCLouvain). NC was supported by the Foundation for Training in Industrial and Agricultural Research (FRIA, FNRS), and SH by the Belgian Federal Public Service of Health, Food Chain Safety and Environment.

## DECLARATION

Conflicts of interest: The authors declare that they have no conflict of interest. Compliance with ethical standards.

## Notes

### Competing Interest Statement

The authors have declared no competing interest.

