## Supplementary Fig S1 & Fig S2 for "Salicaceae as potential host plants of *Xylella fastidiosa* in European temperate regions"

**Fig. S1** Distribution of *X. fastidiosa* KLN 59.3 strain in xylem vessels 9 months after mechanical inoculation in the stem of *P. tremula*, *S. alba* and *P. canescens*. The diagram of each plant part was designed by placing the locations where the bacteria were observed by confocal microscopy on the cross-sections visualized by electron microscopy, as described in part **a**. Red colored circle is for the observation of totally obstructed vessels by the bacterium and green circle for the partially obstructed vessels. The numbers in black on the side of the colored circles represent the number of vessels observed near each other with the same bacterial state (partially obstructed or totally obstructed). In **b.** The diagrams for each individual. The annotation of a plant part without the representation of the section diagram means that no bacteria were observed in the section.

**a**

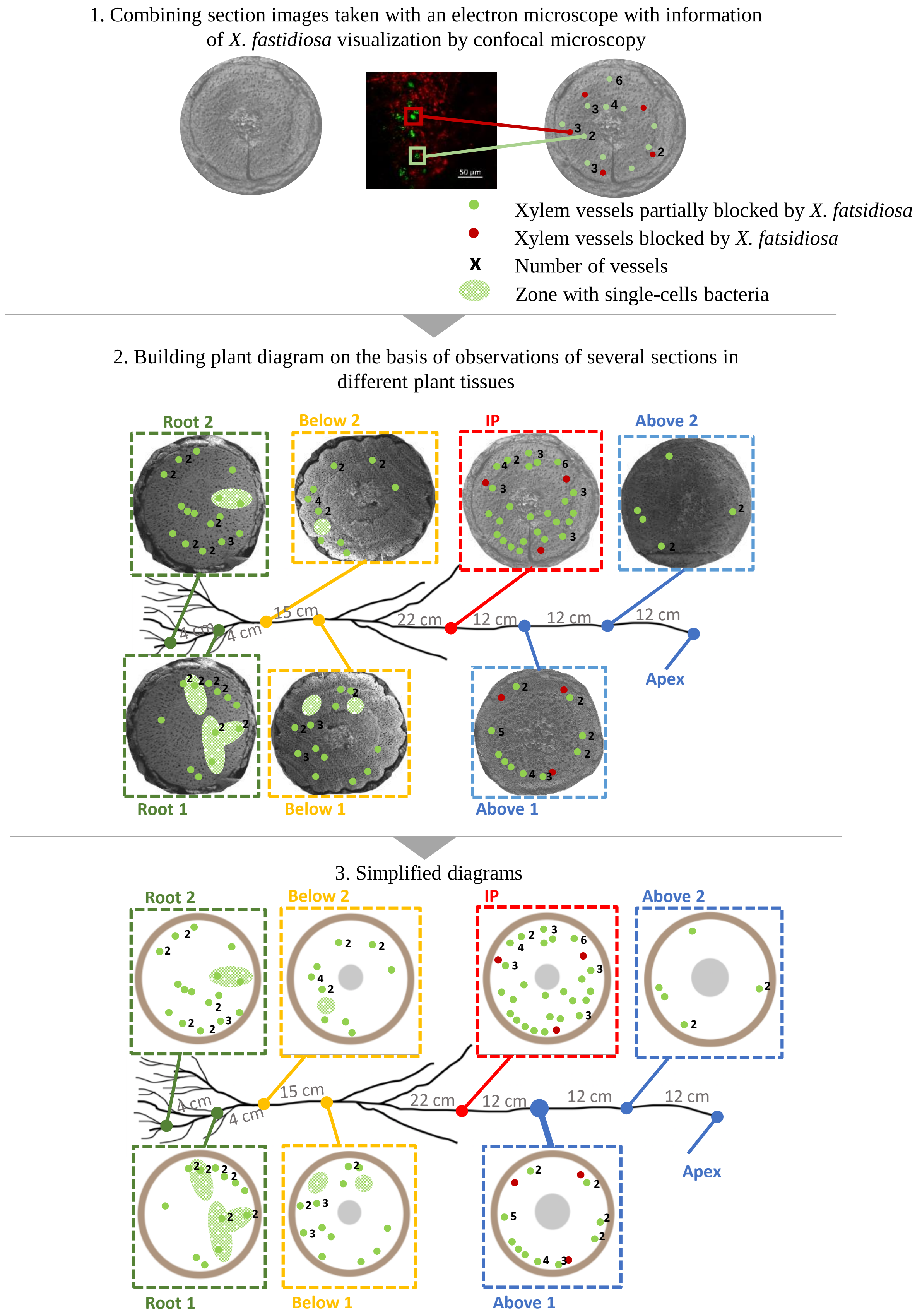

**b**

*Populus tremula* – 22°C growing conditions

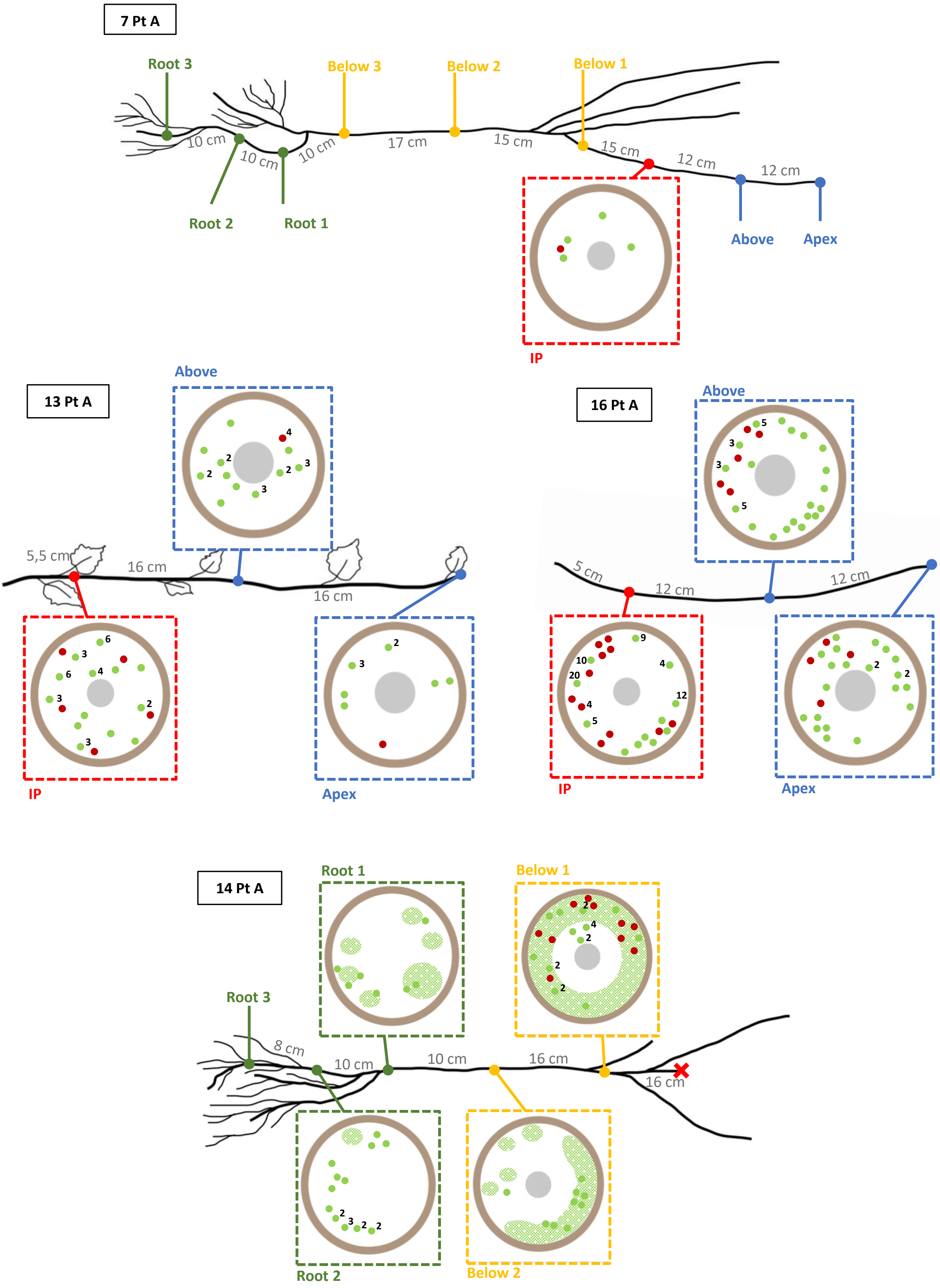

***Populus tremula* – 28°C growing conditions**

**9 Pt B**

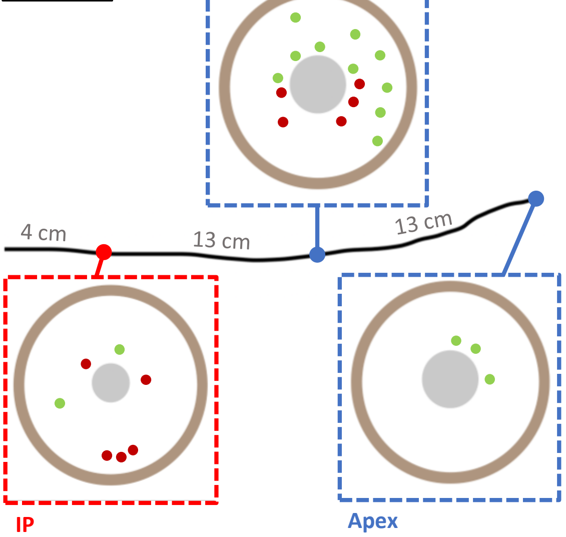


**12 Pt B**

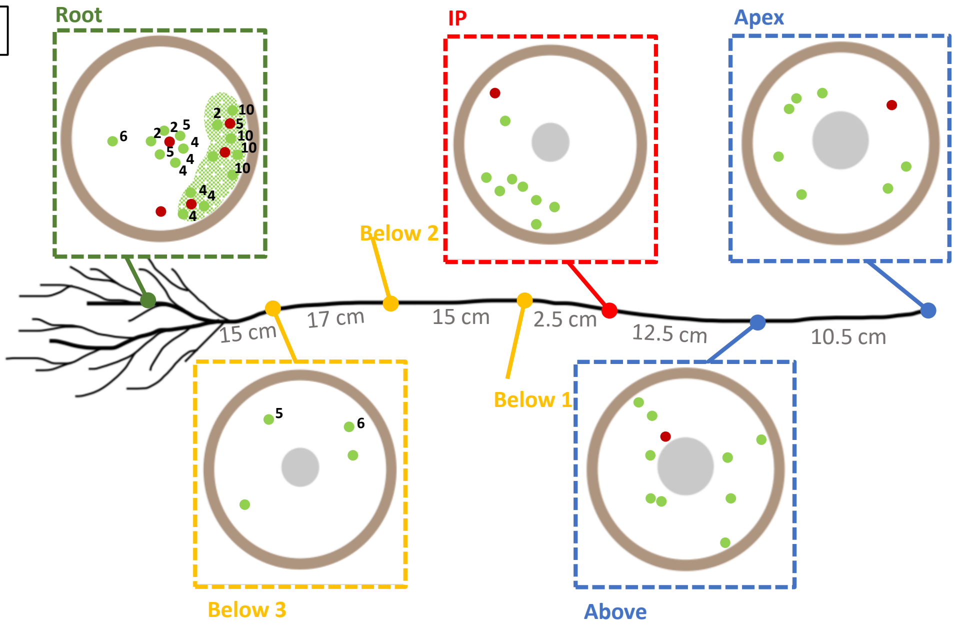


**14 Pt B**

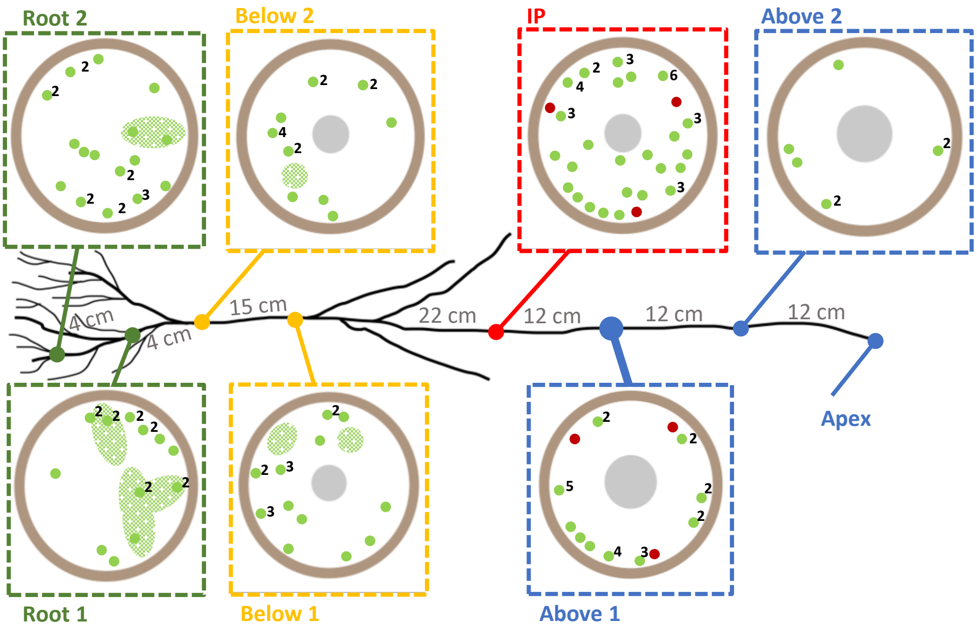

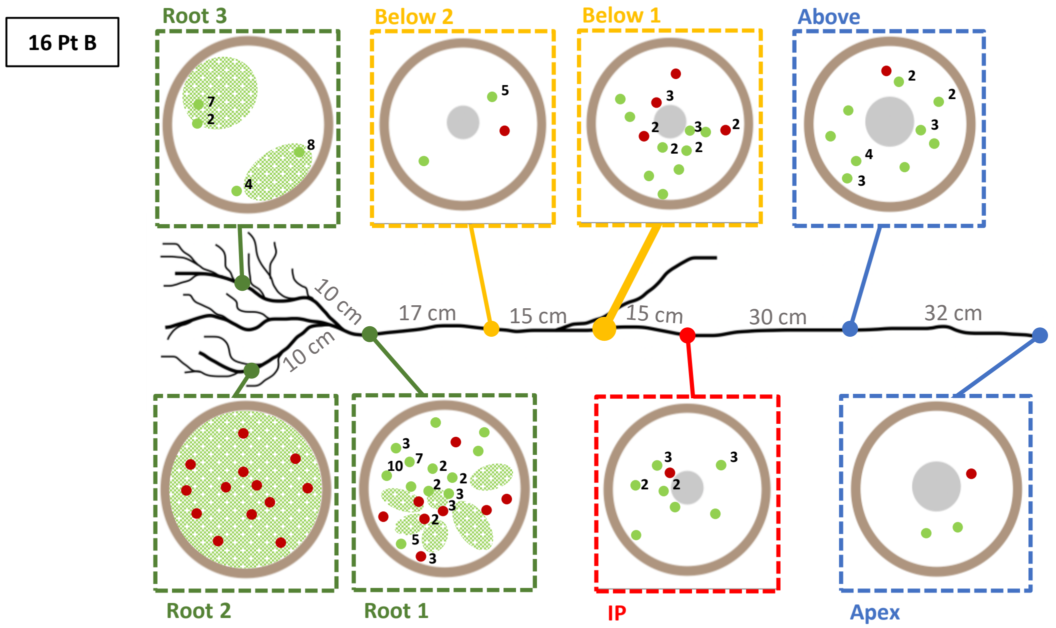


*Salix alba* – 22°C growing conditions

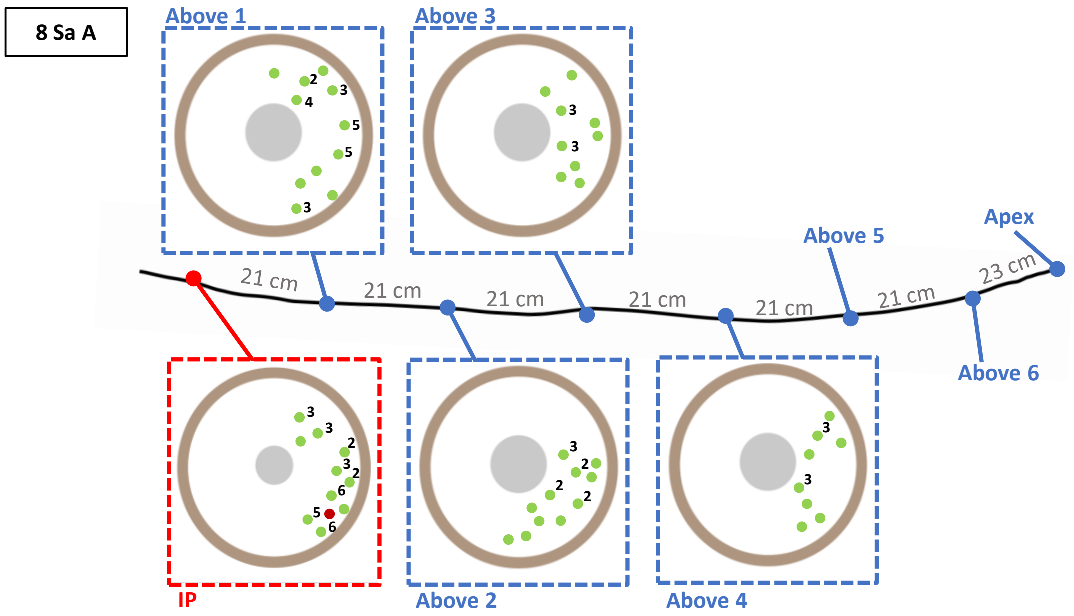


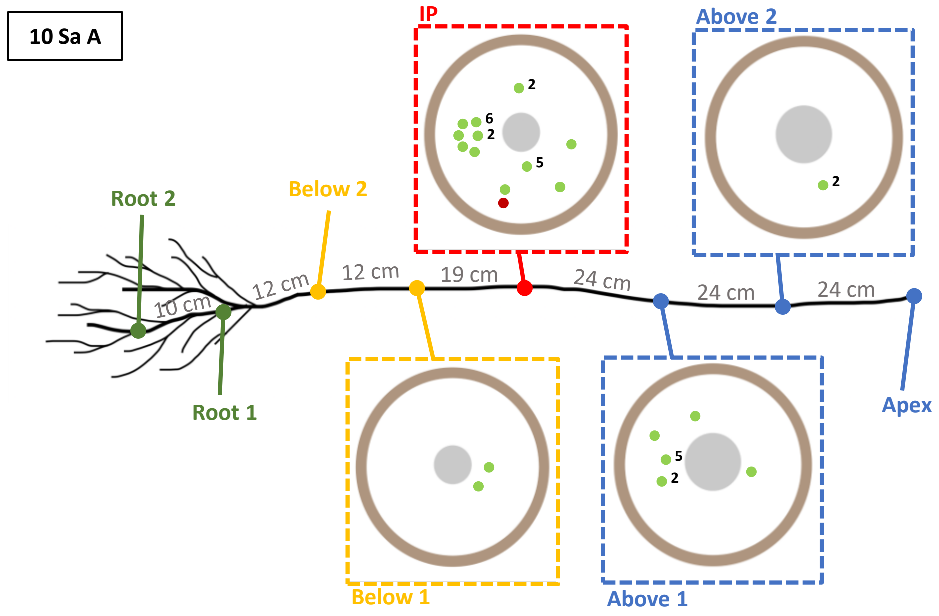

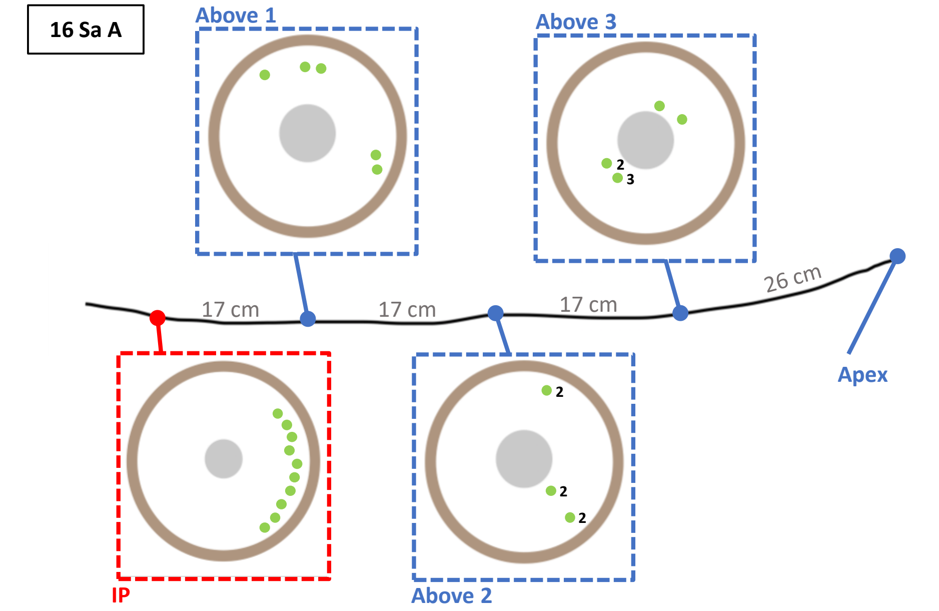


***Salix alba* – 28°C growing conditions**

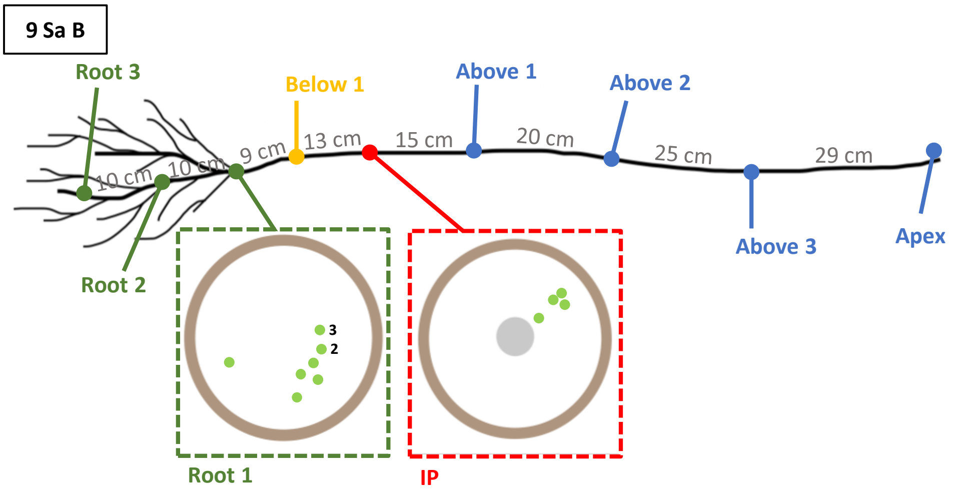


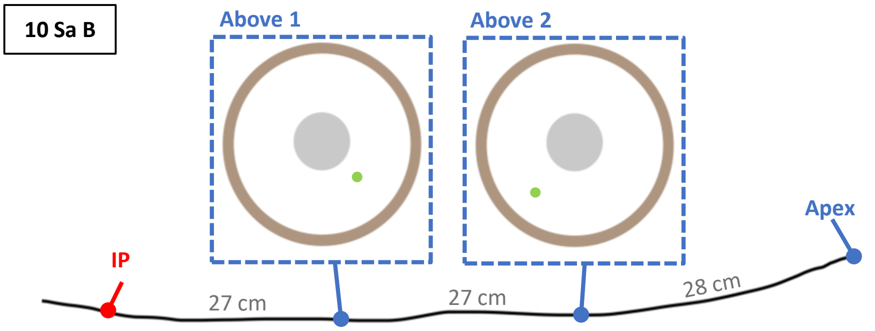


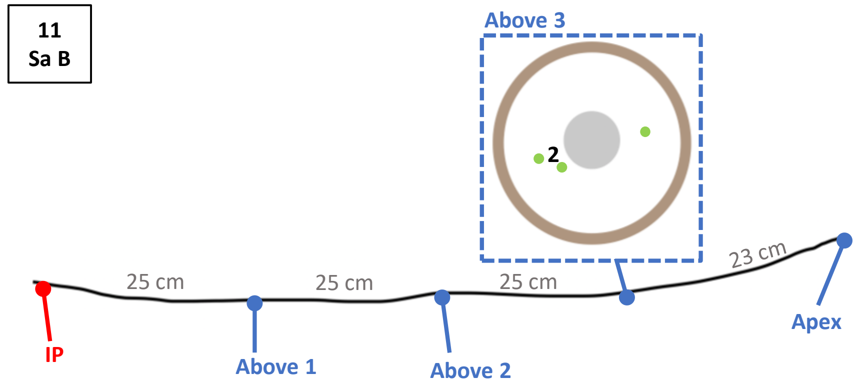

*Populus canescens* – 22°C growing conditions

11 Pc A

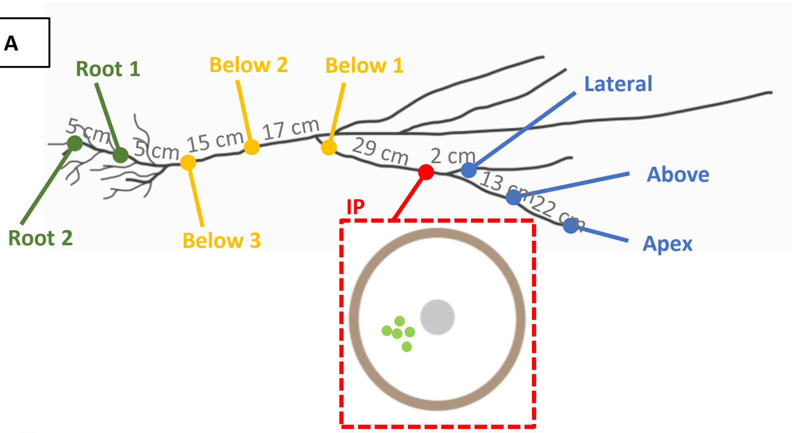


14 Pc A

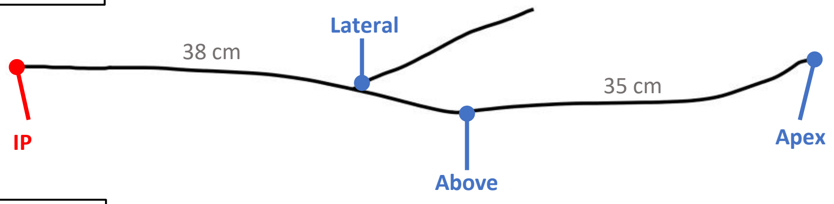


15 Pc A

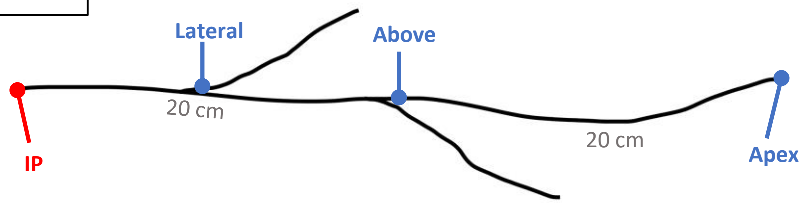


*Populus canescens* – 28°C growing conditions

12 Pc B

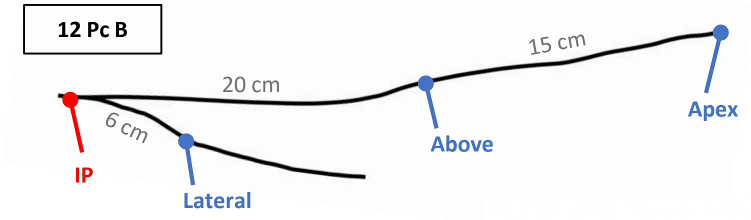


13 Pc B

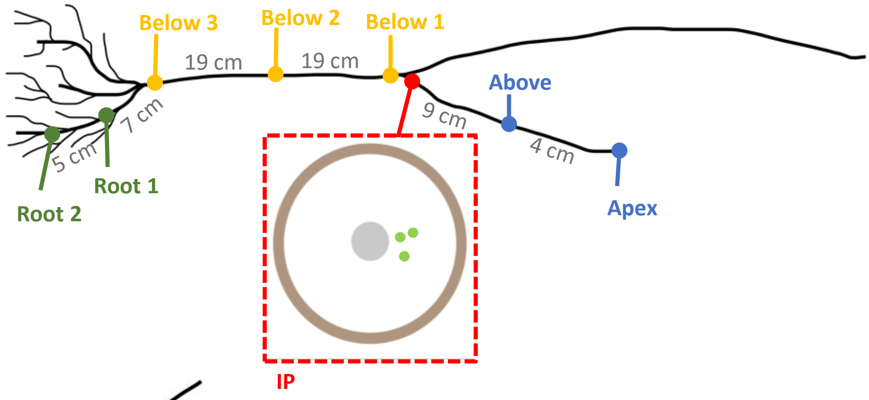


14 Pc B

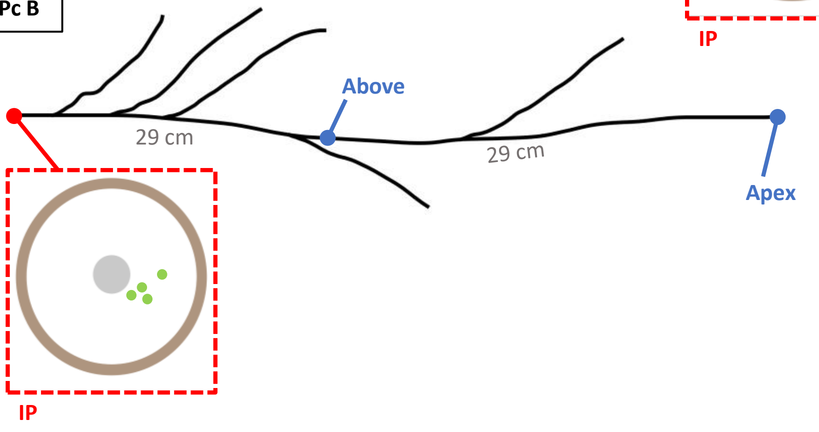

**Fig. S2** *X. fastidiosa* in xylem vessels of one inoculated *P. tremula* plant at the inoculation point. **a.** Diagram representing the sections visualized by confocal and electron microscopy. Red circle is for the observation of totally obstructed vessels by the bacterium and green circle for the partially obstructed vessels. **b**. *X. fastidiosa* depicted in green, expressing the GFP in xylem vessels observed with confocal microscopy at the location of the pink frame in "a". **c**. Visualization of about a part of the cross section of the inoculated stem (at the location of the pink frame in "a") with an electron microscope in LM mode. The white bar represents a 100-µm scale. **d**. Zoom on the xylem vessels containing the bacterial mat, observations with electron microscope at the gentle beam (GB-H) mode. The white bar represents a 1-µm scale. **e.** Zoom on the bacterial biofilm, observations with electron microscope at GB-H mode.

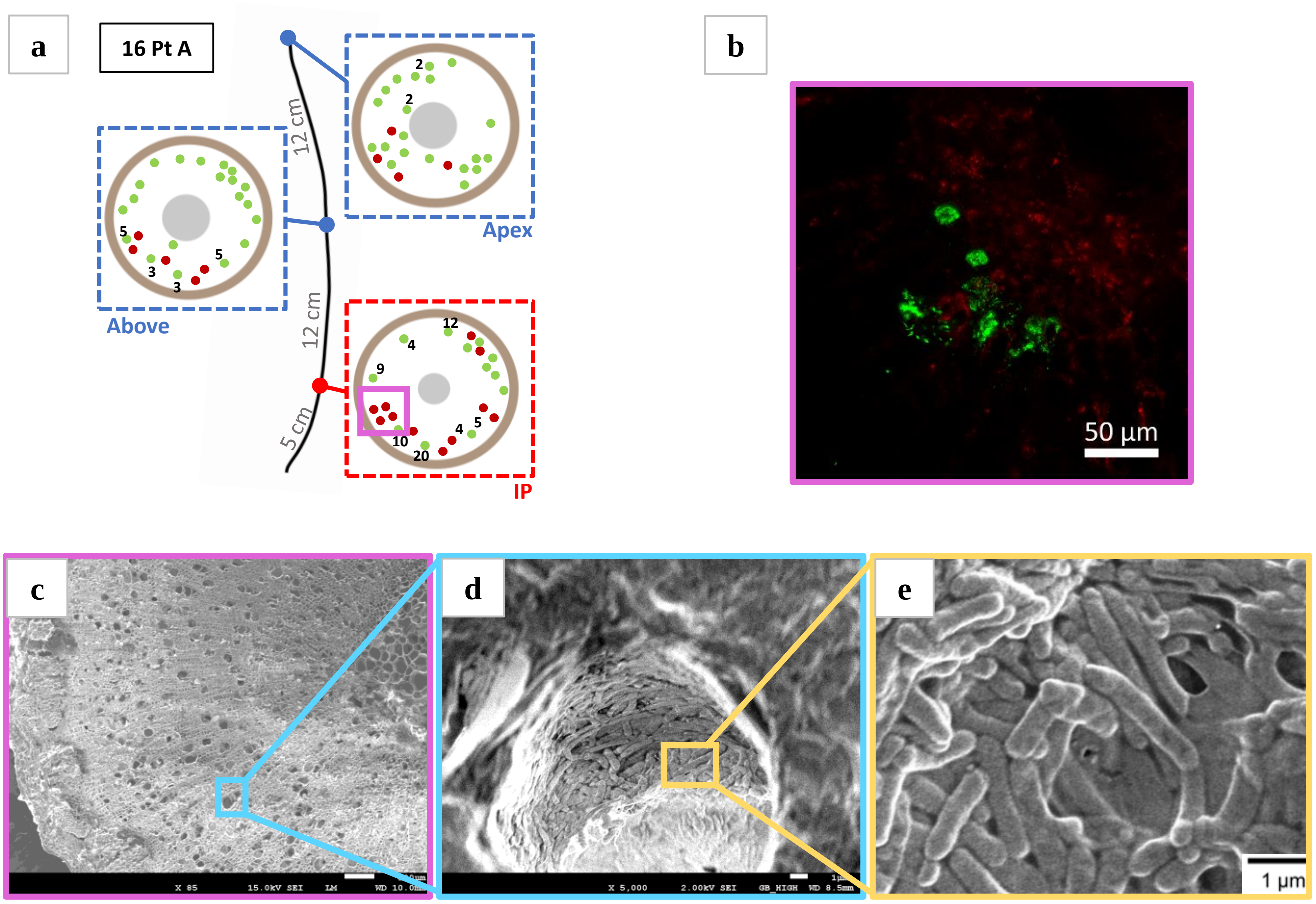
